# Towards an informed choice of diffusion MRI image contrasts for cerebellar segmentation

**DOI:** 10.1101/2025.03.10.642452

**Authors:** Jon Haitz Legarreta, Zhou Lan, Yuqian Chen, Fan Zhang, Edward Yeterian, Nikos Makris, Jarrett Rushmore, Yogesh Rathi, Lauren J. O’Donnell

## Abstract

The fine-grained segmentation of cerebellar structures is an essential step towards supplying increasingly accurate anatomically informed analyses, including, for example, white matter diffusion magnetic resonance imaging (MRI) tractography. Cerebellar tissue segmentation is typically performed on structural magnetic resonance imaging data, such as T1-weighted data, while connectivity between segmented regions is mapped using diffusion MRI tractography data. Small deviations in structural to diffusion MRI data co-registration may negatively impact connectivity analyses. Reliable segmentation of brain tissue performed directly on diffusion MRI data helps to circumvent such inaccuracies. Diffusion MRI enables the computation of many image contrasts, including a variety of tissue microstructure maps. While multiple methods have been proposed for the segmentation of cerebellar structures using diffusion MRI, little attention has been paid to the systematic evaluation of the performance of different available input image contrasts for the segmentation task. In this work, we evaluate and compare the segmentation performance of diffusion MRI-derived contrasts on the cerebellar segmentation task. Specifically, we include spherical mean (diffusion-weighted image average) and b0 (non-diffusion-weighted image average) contrasts, local signal parameterization contrasts (diffusion tensor and kurtosis fit maps), and the structural T1-weighted MRI contrast that is most commonly employed for the task. We train a popular deep-learning architecture using a publicly available dataset (HCP-YA), leveraging cerebellar region labels from the atlas-based SUIT cerebellar segmentation pipeline. By training and testing using many diffusion-MRI-derived image inputs, we find that the spherical mean image computed from b=1000 s/mm^2^ shell data provides stable performance across different metrics and significantly outperforms the tissue microstructure contrasts that are traditionally used in machine learning segmentation methods for diffusion MRI.

**Key points:**

- We provide evidence about the performance of different dMRI contrasts for cerebellar structure segmentation using a deep learning neural network.
- The diffusion MRI spherical mean provides improved and stable cerebellar structure segmentation performance.
- The spherical mean is easy to compute and can be used for cerebellar structure segmentation on retrospective clinical diffusion MRI data.

## 1. Introduction

The cerebellum is a large brain region attached to the brainstem that hosts a large proportion of the neurons in the human brain. It plays a critical role in movement, cognition, language, emotion, and perception (Baumann et al., 2015; Buckner, 2013; Murdoch, 2010; Riedel et al., 2015; Strick et al., 2009), and thus, it is one of the most important structures in the brain. Yet, the analysis of cerebellar neuroanatomy on *in vivo* magnetic resonance imaging (MRI) data is challenging due to several factors (Lundell & Steele, 2024), including acquisition aspects due to its location (e.g. the whole extension of the cerebellum may not be included in the field of view of MRI acquisitions whose focus is not the cerebellum) or a limited resolution, its relatively small size (it represents only 10% of the weight of the brain (Azevedo et al., 2009; Herculano-Houzel, 2009)) or its strikingly complex anatomy, including its tightly foliated cerebellar cortex. Thus, the availability of cerebellar data and the corresponding annotations is impacted, and most available MRI segmentation work has been focused in developing methods that are demonstrated on the cerebrum, with a less detailed parcellation of the cerebellar structures.

The cerebellum is composed of a superficial three-layered cortex and underlying cerebellar white matter in which the deep cerebellar nuclei (DCN) are located. The cerebellar cortex is generally divided into ten distinct lobules (denoted by Roman numerals: I-X) by transverse fissures. In addition, the cortex can be divided into left and right hemispheres, which are separated from a midline region called the vermis by paired longitudinal sulci. Regions of the cerebellar cortex are thus named based on the hemisphere (L for left, R for right, or vermis) and their associated lobule (I-X). In addition, the hemispheres of lobule VII are often subdivided into two regions referred to as crus I and II. The deep cerebellar nuclei are comprised of four paired nuclei: the dentate, globose, emboliform and fastigial. The dentate is the largest and most lateral of the DCN. The fastigial nucleus is located medially, close to the mid-sagittal plane. Finally, the globose and emboliform nuclei (often together referred to as the interposed nuclei) are located between the dentate and fastigial nuclei. See (Altman & Bayer, 1996) for a detailed description of the human cerebellar anatomy.

Accurate delineation of such cerebellar structures from MRI data is an essential step towards the goal of studying the cerebellar neuroanatomy and its connectivity pattern *in vivo* and non-invasively. Although numerous automatic methods have been proposed for cerebellar structure segmentation, many of these rely on structural MRI (sMRI) data, such as T1-weighted (T1w) or T2-weighted (T2w) acquisitions, or segment only part of the cerebellar structures (see recent examples of such works in (Beliveau et al., 2021; Carass et al., 2018; Faber et al., 2022; Han et al., 2020; Morell-Ortega et al., 2024). In order to study brain connectivity patterns from tractography-derived data, such segmentation maps need to be warped to the diffusion MRI (dMRI) volumes, which may result in errors from the registration process that may negatively impact the study of connectivity (Cheng et al., 2020; Theaud et al., 2022; F. Zhang et al., 2024).

Performing brain tissue segmentation directly from dMRI features is a strategy that avoids the need for inter-modality registration and can improve the anatomical specificity of tractography and derived connectivity analyses (Theaud et al., 2022; F. Zhang et al., 2021). Deep learning methods that have been recently proposed for dMRI segmentation have employed model-based (i.e. based on frameworks representing the diffusion process) tissue microstructure contrasts derived from diffusion tensor imaging (DTI) and diffusion kurtosis imaging (DKI). Summarized in Table 1, these contrasts (or a combination of them) have been shown to enable successful brain structure segmentation in the literature. A few of these studies have used diffusion MRI data to train a neural network to automatically delineate a subset of the cerebellar structures. Gaviraghi et al. (Gaviraghi et al., 2021) segmented the dentate nuclei on b0 dMRI images using regular convolutional classification neural networks. Kim et al. (Kim et al., 2020) proposed a method to segment DCN on high-field (7T) b0 images using a semi-supervised neural network model, but excluded the fastigial nuclei due to the difficulty in segmenting them. Bermudez Noguera et al. (Noguera et al., 2019) segmented the dentate nuclei using a U-Net and a multi-contrast (T1w, T2w, FA) approach.

**Table 1.** Summary of dMRI-based features used for brain tissue segmentation. Legend: AFD: apparent fiber density; FA: fractional anisotropy; MD: mean diffusivity; RD: radial diffusivity; DT: diffusion tensor; MK: mean kurtosis; AK: axial kurtosis; RK: radial kurtosis; FOD: fiber orientation density; DWI: diffusion-weighted image data. CSF: cerebrospinal fluid; GM: gray matter; WM: white matter.

| Model-based | dMRI contrast | Derived from | Segmented tissue | Deep learning | Citations |
| --- | --- | --- | --- | --- | --- |
| Yes | FA, eigenvalues, AK, MK, RK | DTI, DKI | CSF, GM, WM | Yes | (F. Zhang et al., 2021) |
| Yes | AFD, AD, MD RD | DTI, FOD | Multiple (10 brain regions) | Yes | (Theaud et al., 2022) |
| Yes | FA, MD, eigenvalues | DTI | Multiple (101 brain regions) | Yes | (F. Zhang et al., 2024) |
| Yes/No | spherical mean, FA, MD | DWI, DTI | CSF, GM, WM | No | (Ciritsis et al., 2018) |
| No | b0 image, spherical mean maps | DWI | CSF, GM, WM | No | (Cheng et al., 2020; Sun et al., 2019; J. Wang et al., 2020) |
| No | b0 image | DWI | Cerebellar nuclei | Yes | (Gaviraghi et al., 2021; Kim et al., 2020) |
| No | b0 image | DWI | Dentate nuclei | Yes | (Gaviraghi et al., 2021; Kim et al., 2020) |
| No | Multiple contrasts investigated | DWI, DTI, DKI investigated | Multiple (34 cerebellar regions) | Yes | Ours |

However, there is a lack of information about the relative performance of these different image contrasts for cerebellar tissue segmentation using dMRI data. Furthermore, although the value of model-based microstructure measures has been demonstrated, basic contrasts that can be derived more directly from the raw diffusion-weighted image (DWI) data^1^, such as the spherical mean (average diffusion-weighted) images, have been understudied in the context of deep learning methods, as shown in Table 1.

Thus, despite these efforts, a study on the potential contribution of dMRI-specific contrasts for the cerebellar segmentation task is missing. Additionally, the use of the spherical mean has not yet been considered to automatically segment cerebellar structures. In this work, we provide evidence about the performance of different dMRI contrasts for the segmentation of cerebellar lobules and nuclei using a deep learning neural network. This work may provide useful insight when developing specific methods to improve existing segmentation models.

## 2. Materials and methods

Our overall approach is to evaluate and compare the performance of diffusion-MRI-derived inputs, including raw diffusion-weighted image and local diffusion model data (tensor and kurtosis-derived fit maps), to parcellate cerebellar structures. We adopt an existing deep learning-based (DL) model to demonstrate the performance of such contrasts separately.

### 2.1. Neural network architecture

We adopt the SegResNet backbone proposed by (Myronenko, 2019) to accomplish the cerebellar segmentation task. SegResNet ranked among the top performers at the BraTS 2021 MRI brain tumor segmentation challenge^2^. For this work we use the implementation available in MONAI (Creators MONAI Consortium, n.d.), and set the network to use 3D volumetric data as the input. Details on the experimental settings are provided in the 2.5. Experiments section, and the Appendix section includes a specification about the network architecture. A separate model is trained on each of the considered input image contrasts, including T1w data and dMRI-derived maps (see section 2.4. Input dMRI-derived contrasts).

### 2.2. Datasets

Data from the S1200 release of the Human Connectome Project (HCP) Healthy Young Adult (YA) dataset (Glasser et al., 2013; Sotiropoulos et al., 2013; Van Essen et al., 2013) are used in this work. The HCP-YA dataset features dMRI data acquired at 1.25 mm isotropic resolution for 1000, 2000 and 3000 s/mm^2^ b-value shells, acquired along 270 diffusion weighting directions, plus 18 b=0 s/mm^2^ (b0) acquisitions. A T1w structural MRI scan is available for each participant. The minimally pre-processed dMRI and T1w data at the dMRI 1.25 mm isotropic resolution as released by the dataset authors are used in this work. All data are padded to a fixed size of 192x192x192 voxels in order to facilitate pipeline operation.

### 2.3. Segmentation labels

The SUIT toolbox (Diedrichsen, 2006), which provides cerebellar lobule and DCN segmentations, is used as the silver standard^3^ to provide the labels to train the models. SUIT is an openly available atlas-based human cerebellar analysis tool that has been adopted by scientists for automatic cerebellar region segmentation (see, for example, (Carass et al., 2018; Faber et al., 2022)). The toolbox is run on the T1w participant data using the MATLAB 2020b and SPM 12 releases. We use the segmentations obtained to train our deep neural network models and to measure the model performance. Figure 1 shows a volumetric depiction of the structures and major label groups. For a complete set of labels, see the Appendix.

**Figure 1.**
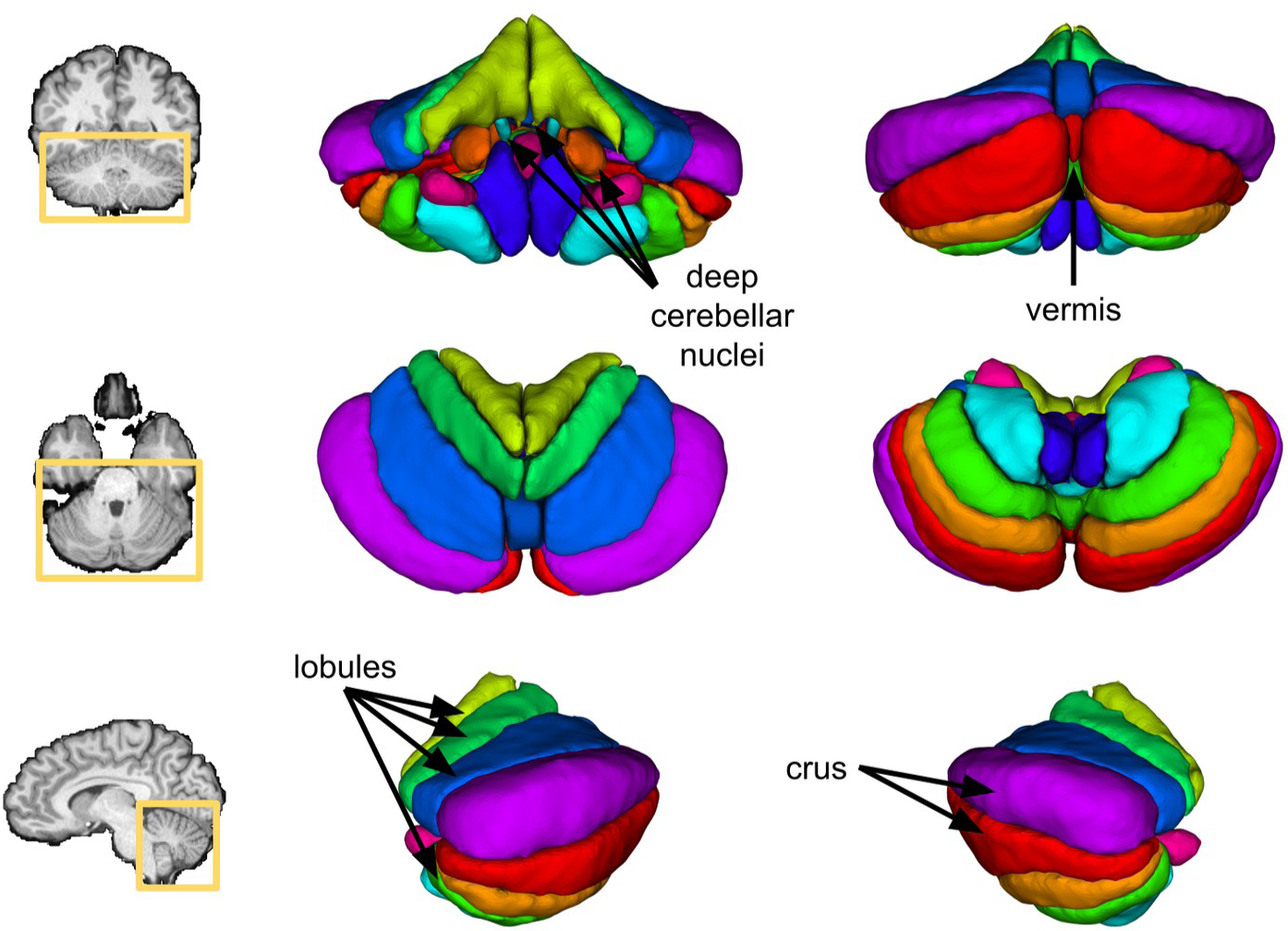
The cerebellum and the cerebellar label groups. Top row: coronal anterior image slice, and anterior and posterior volumetric views of the cerebellar structures; middle row: axial superior image slice, and superior and inferior views; bottom row: sagittal left image slice, and left and right views.

### 2.4. Input dMRI-derived contrasts

The input dMRI-derived data for the neural network model are computed from DWI data and diffusion tensor and diffusion kurtosis model fits. From the DTI model, we include the eigenvalues of the tensor (E1, E2, E3), the mean diffusivity (MD), the radial diffusivity (RD), and the fractional anisotropy (FA); from the DKI model, we include the axial kurtosis (AK), the mean kurtosis (MK), and the radial kurtosis (RK). These contrasts have been shown in the literature to enable successful brain structure segmentation (Table 1). We also include the mean b=0 s/mm^2^ or baseline image (b0), as well as the diffusion-weighted image averaged across all gradient encoding directions (spherical mean, SM) for each b-value (SM1k, SM2k, and SM3k for the b=1000 s/mm ^2^, b=2000 s/mm^2^, and b=3000 s/mm^2^ shells, respectively) and for all b-values (SM). We compute these contrasts using DIPY (Garyfallidis et al., 2014) and SCILPY (https://github.com/scilus/scilpy), as well as in-house software. DKI-derived contrasts are computed using all three shells following the recommended b ≤ 3/(D*K) b-value upper bound and assuming typical values of D ≈ 1e-3 mm2/s and K ≈ 1 provided by (Jensen & Helpern, 2010). We also include T1w image data, as T1w is the primary MRI contrast used for segmentation tasks and provides a benchmarking reference for dMRI-based contrasts. Table 2 summarizes the input image contrasts considered in this work. Figure 2 shows coronal anterior close-up views of the cerebellar region across the input image contrasts on a given participant. The deep cerebellar nuclei (DCN) are highlighted (yellow arrows) in one of the hemispheres for those contrasts where they are most apparent.

**Figure 2.**
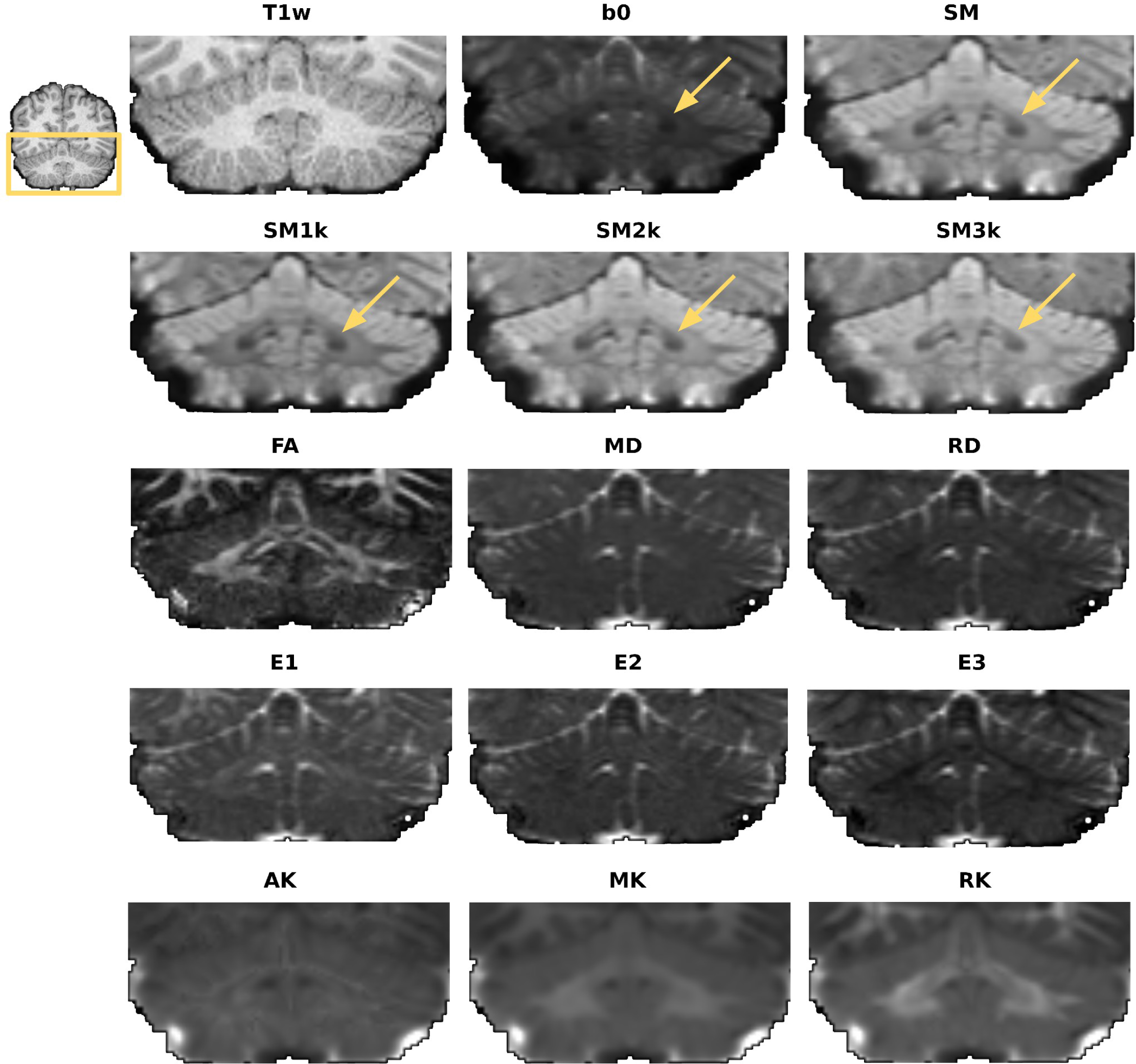
Close-up view of the cerebellum across the considered contrasts. Yellow arrows point to the DCN structure area. All coronal anterior views.

**Table 2.**
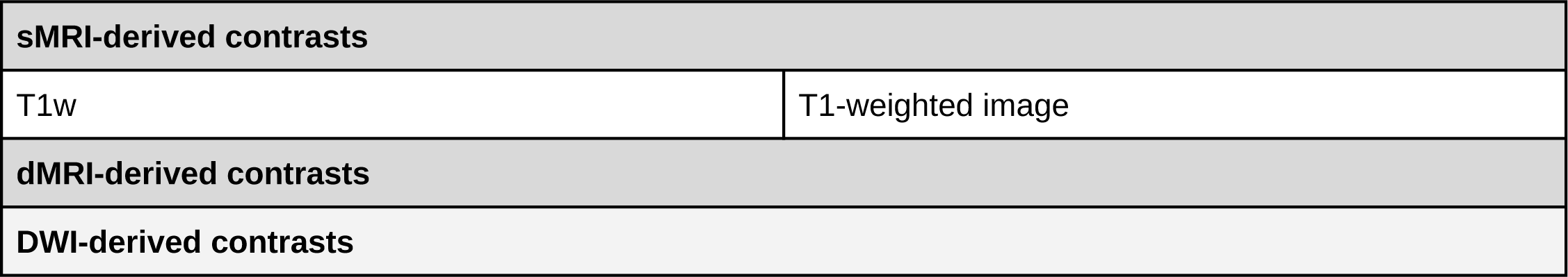
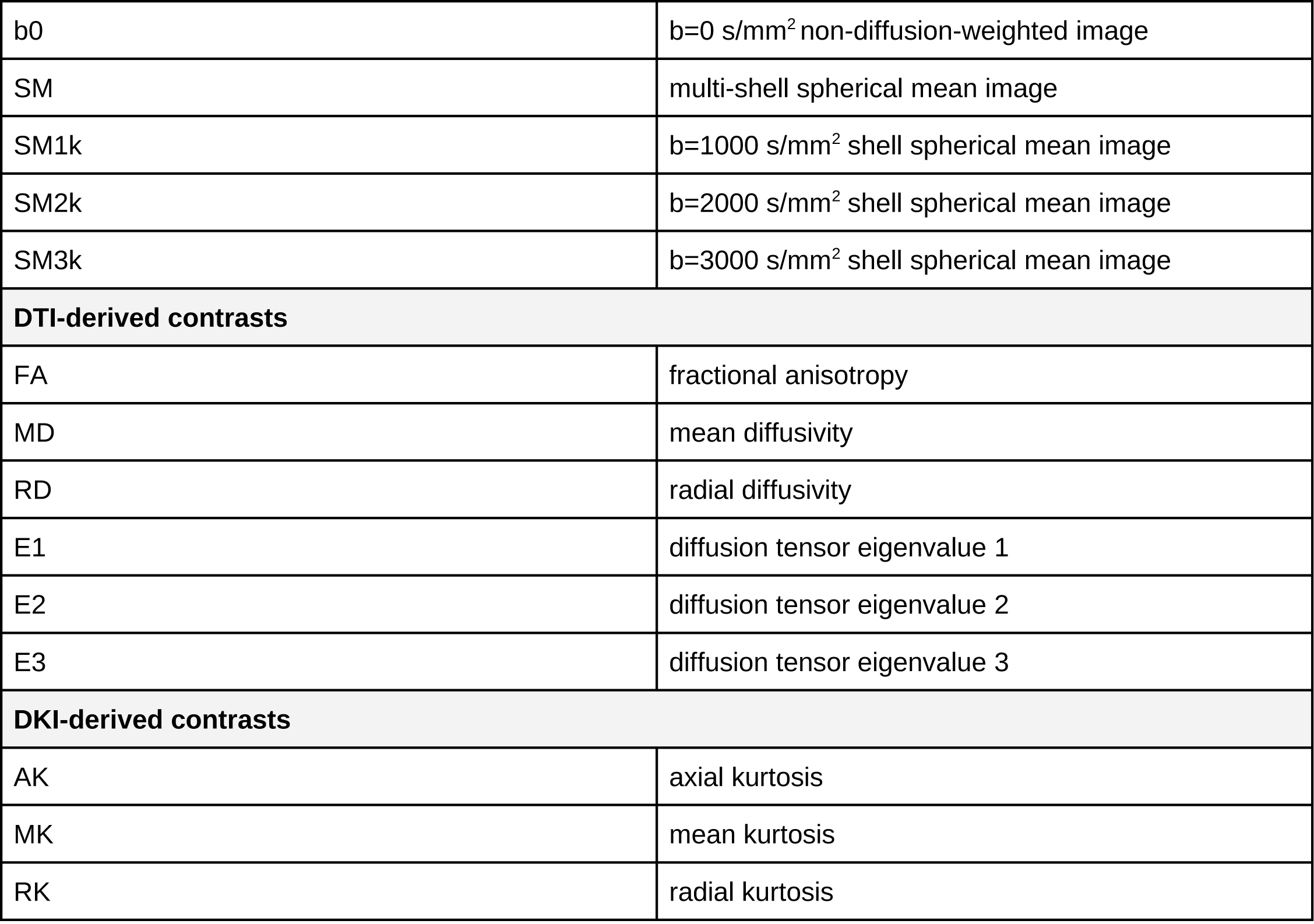
Image contrasts considered in this work as the input of the deep neural network model.

### 2.5. Experiments

We use data from 100 participants from the HCP-YA S1200 release to train and test the models. All data, including labels, were quality checked visually, and participants showing incomplete coverage of the cerebellum, large distortions, or evident mislabelling were excluded to build the set of 100 participants. We train the deep learning models separately on the T1w data and on each of the contrasts derived from the dMRI data. Five-fold cross validation is used and results are reported on the average performance across folds. The data are split into training and test sets according to the 80/20 rule; 20 participants are used for validation purposes in each training set.

The batch size is set to 1 to accommodate the network parameters and input data size into the GPU memory constraints. Image intensities are scaled to the [0,1] range, and random flips with p=0.5 probability along the first axis are used for data augmentation following the practices suggested for the chosen neural network (Myronenko, 2019). The optimization framework used is the same across all models so that the performance could be attributed only to the contrast modality: the deep learning model is trained using the popular Adam optimizer (Kingma & Ba, 2015), the initial learning rate being set to 1e-4 following (Myronenko, 2019), and a learning rate decay scheduler with a factor of 0.5, patience 10 and a lower bound of 1e-8. It was experimentally verified that larger learning rates would not improve the achieved final validation metric, and a sustained value of 1e-4 would provide reasonable training regimes before plateauing. The models are trained for 250 epochs, fixed after experimental evidence showing appropriate model convergence across contrasts. The models are trained to minimize the unweighted sum of the cross-entropy and Dice losses. It has been shown (Taghanaki et al., 2019) that this combined loss outperforms the results obtained by each separately, and has become a popular choice for segmentation. The validation metric used for model selection is the Dice similarity coefficient (DSC) (Dice, 1945).

In order to assess the contribution of the different dMRI contrasts for the deep learning-based cerebellar segmentation task, the following experiments are performed:

#### 2.5.1. Experiment 1: Comparison of performance across dMRI images and T1w data

This experiment was designed to quantify and compare the segmentation performance obtained from using different dMRI contrasts and T1w data. The deep neural network model is trained and tested separately on each input image contrast listed in Table 2 on the cerebellar structure segmentation task, and evaluation metrics (section 2.6. Evaluation metrics) are computed to assess model performance.

#### 2.5.2. Experiment 2: Generalization of models to fewer gradient directions

We explore the ability of the diffusion-MRI derived images to provide with sufficient contrast input information to the deep neural network to accomplish the cerebellar segmentation task when using fewer diffusion gradient-encoding directions. The data are subsampled across the gradient directions to keep 60, 30 and 20 directions in each shell. For each set of directions and shells, the diffusion data corresponding to the closest gradient vectors for optimal uniform coverage are kept following (Caruyer et al., 2013). Note that the subsampled data have lower angular resolution, but maintain the same spatial resolution of the original data. 20 directions represent a common lower bound for DTI-like clinical acquisitions, 60 being a desirable value for HARDI acquisitions, and 30 being an intermediate choice between the two schemes. See for example (Descoteaux, 2015; Tournier et al., 2013) for considerations on DTI-like and HARDI acquisitions. New input contrast data are then generated from the subsampled data for the two top-performing contrasts from Experiment 1. The neural network models trained on the corresponding complete set of directions for each contrast and direction set are employed to segment the cerebellar regions on the newly computed, subsampled input data.

### 2.6. Evaluation metrics

Following (Maier-Hein et al., 2024) we include overlap-, boundary-, and property-based metrics in our evaluation. The overlap metric used in this work is the DSC, which measures the degree of spatial overlap between two structures. Boundary-based metrics include the 95% percentile Hausdorff distance (HD95), the mean surface distance (MSD), and the center of mass distance (CMD). HD95 measures the 95th percentile of the maximum distance of the predicted segmentation to the nearest point in the silver standard, MSD represents the mean value across the surface distances between the prediction and the silver standard boundaries, and CMD is computed as the average distance across the structures of interest of the mass centers (centroids) between the prediction and the silver standard labels. Property-based metrics comprise the volume similarity (VS), which provides a normalized measure of the volume differences between the predicted and silver standard structures (Keim, 1999). Finally, we compute the label detection rate (LDR) as the ratio of the correctly detected number of labels over the silver standard labels. These metrics are computed using the seg-metrics (Jia et al., 2024) and SciPy (Virtanen et al., 2020) Python packages and in-house software.

### 2.7. Statistical analysis

For each metric, a repeated measures ANOVA (rmANOVA) analysis and a pairwise Student’s t-test corrected for multiple comparisons (using the false discovery rate (FDR) procedure (Benjamini & Hochberg, 1995)) are carried out to determine whether the observed differences were significant, and to provide statistical significance values. For each metric, the pairwise significance is computed against the top performing contrast. In this work, probability values above p > 0.05 are considered non-significant; the significance levels are noted as follows: (*) indicates p < 0.05, (**) indicates p < 0.01, and (***) indicates p < 0.001.

## 3. Results

Tables 3, 4, 5, 6, 7 and 8 show the results for each metric and contrast. In each table, average results across participants are shown (standard deviation values are shown in parentheses). The values in each column are computed considering the segmentation labels corresponding to the cerebellar structures of interest (described in 1. Introduction). The “all” column includes all considered structures. Boldface text is used to denote the top-performing values across all relevant structures; statistical significance is indicated in parentheses following the usual notation mentioned in section 2.7. Statistical analysis.

**Table 3.**
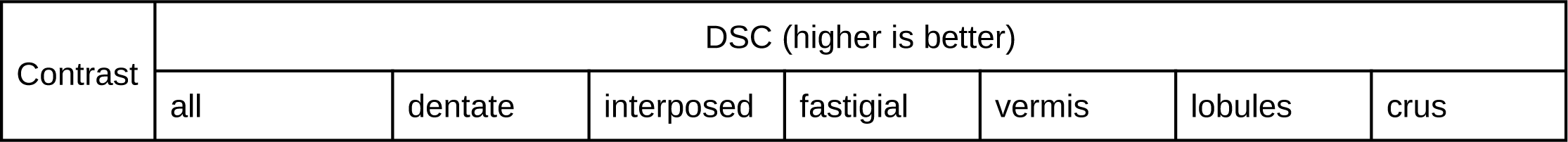
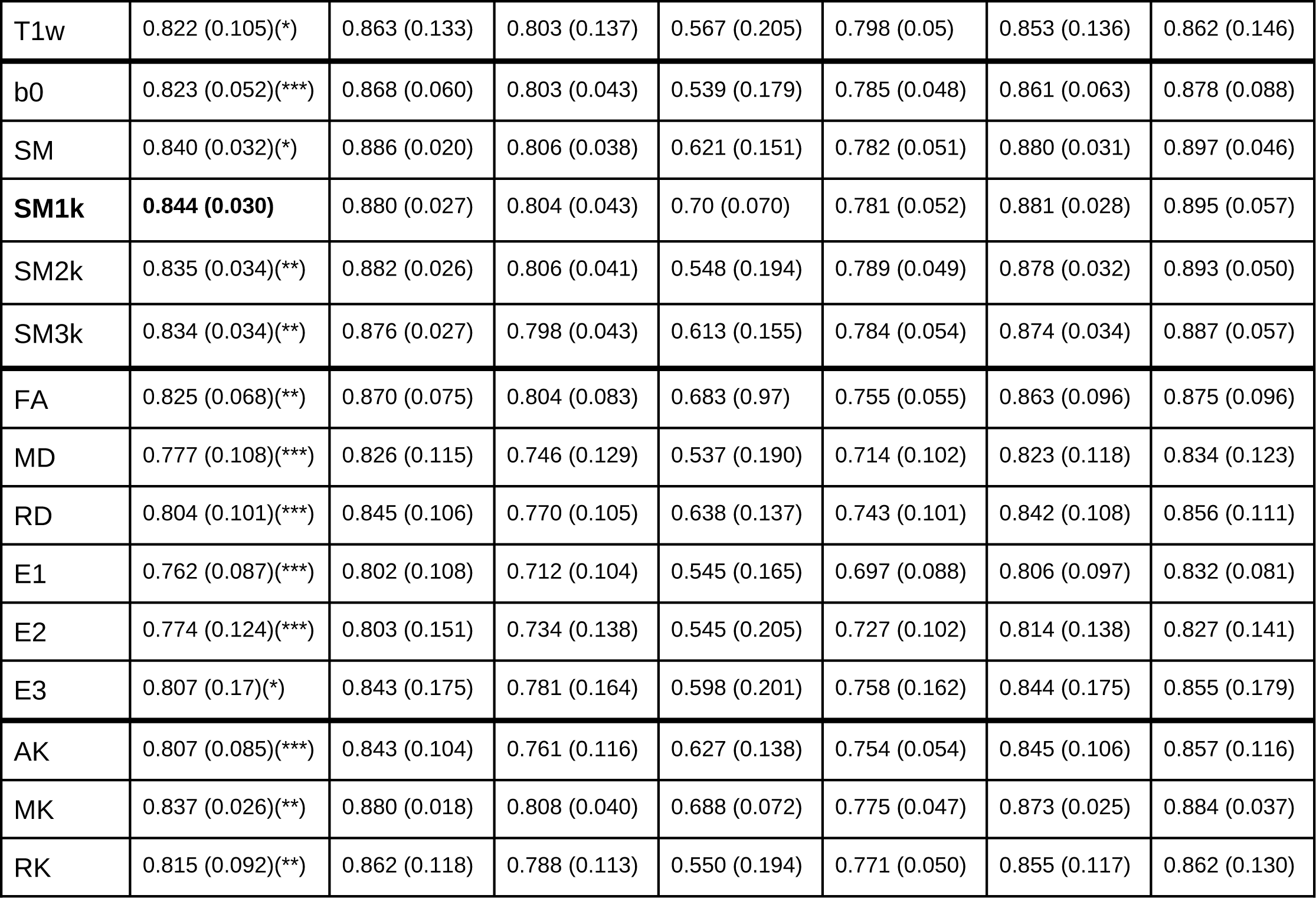
DSC performance across participants. Mean (standard deviation) values across participants for different target structures. Best mean values are highlighted in bold. Asterisks indicate contrasts with significantly worse performance than the top performing contrast.

**Table 4.** HD95 performance across participants. Mean (standard deviation) values across participants for different target structures. Best mean values are highlighted in bold. Asterisks indicate contrasts with significantly worse performance than the top performing contrast.

| Contrast | HD95 (lower is better) |  |  |  |  |  |  |
| --- | --- | --- | --- | --- | --- | --- | --- |
|  | all | dentate | interposed | fastigial | vermis | lobules | crus |
| T1w | 2.801 (4.671)(*) | 2.622 (4.889) | 1.947 (2.486) | 1.363 (0.395) | 1.440 (0.433) | 3.241 (6.162) | 4.784 (10.387) |
| b0 | 2.056 (2.773) | 1.716 (2.60) | 1.359 (0.739) | 1.338 (0.159) | 1.386 (0.195) | 2.171 (3.740) | 3.715 (7.670) |
| **SM** | **1.443 (0.911)** | 1.259 (0.058) | 1.260 (0.051) | 1.321 (0.154) | 1.422 (0.401) | 1.435 (1.014) | 1.760 (2.984) |
| SM1k | 1.446 (0.979) | 1.297 (0.129) | 1.260 (0.051) | 1.293 (0.117) | 1.298 (0.176) | 1.448 (1.285) | 1.776 (3.081) |
| SM2k | 1.553 (1.230) | 1.357 (1.005) | 1.257 (0.040) | 1.323 (0.161) | 1.389 (0.198) | 1.618 (1.589) | 1.943 (3.639) |
| SM3k | 1.667 (1.654) | 1.281 (0.106) | 1.264 (0.063) | 1.316 (0.132) | 1.393 (0.203) | 1.807 (2.414) | 2.214 (4.558) |
| FA | 1.956 (3.147) | 1.938 (3.612) | 1.465 (1.475) | 1.368 (0.394) | 1.427 (0.227) | 2.049 (4.120) | 3.205 (7.931) |
| MD | 3.096 (5.112)(*) | 2.359 (3.661) | 1.795 (1.908) | 1.490 (0.392) | 1.578 (0.338) | 3.516 (6.047) | 4.763 (8.531) |
| RD | 3.267 (11.481) | 2.939 (11.831) | 2.552 (10.627) | 2.703 (11.058) | 2.524 (10.201) | 3.528 (11.891) | 4.456 (13.530) |
| E1 | 3.569 (7.891)(*) | 3.556 (11.227) | 2.239 (5.432) | 1.725 (0.629) | 2.604 (8.720) | 4.062 (8.871) | 5.046 (8.258) |
| E2 | 4.267 (11.664) | 3.820 (9.821) | 2.969 (9.863) | 2.624 (9.797) | 2.594 (10.382) | 4.796 (12.642) | 7.012 (15.787) |
| E3 | 4.958 (16.914) | 4.609 (16.210) | 1.837 (5.751) | 2.105 (7.673) | 4.416 (15.613) | 5.183 (17.706) | 5.173 (17.165) |
| AK | 2.236 (3.476) | 2.202 (3.912) | 1.583 (1.693) | 1.470 (0.433) | 1.469 (0.253) | 2.494 (4.632) | 3.412 (7.720) |
| MK | 1.597 (0.982) | 1.331 (0.755) | 1.299 (0.443) | 1.311 (0.131) | 1.402 (0.192) | 1.632 (1.267) | 2.267 (3.384) |
| RK | 2.208 (3.877) | 2.062 (4.182) | 1.590 (1.922) | 1.383 (0.542) | 1.432 (0.257) | 2.394 (5.0) | 3.638 (8.794) |

### 3.1. Experiment 1: Comparison of performance across dMRI images and T1w data

In this experiment, we first provide a quantitative analysis of the results for each metric, followed by an overall ranking of all input image contrasts. We then provide a qualitative analysis of example cerebellar segmentation visualizations resulting from the employed image contrasts.

#### 3.1.1. Quantitative analysis

The Dice similarity coefficient (DSC) metric evidence in Table 3 shows that the neural network model performs best when the spherical mean based contrast data (i.e., SM, SM1k, SM2k, SM3k) are used as the input data. Among the spherical mean contrasts, SM1k provides the best DSC. This superior performance is consistent across labels, with SM1k being notably better at delineating the fastigial nuclei. Among the DTI- and DKI-derived maps, MK provides the best results, with E1 and E2 providing the least competitive results. It is also verified that performances are impacted when the smaller structures (interposed, fastigial, vermis) are considered. The post hoc statistical analysis shows that SM1k provides significantly higher performance compared to all other contrasts.

The 95% percentile Hausdorff distance (HD95) metric results are shown in Table 4. Results show that the various spherical mean input data (SM, SM1k, SM2k, SM3k) achieve good performance, with the use of the DKI-derived MK contrast comparing slightly better than SM3k. The model predictions on the DTI-derived eigenvalues and diffusivity-based (MD and RD) input data provide larger errors compared to the rest of the contrasts. Larger (worse) HD95 values are seen on the crus, probably because it contains the largest structures, and thus segmentation inaccuracies accumulate across its boundaries to a larger extent. Most observed differences are not statistically significant according to the post hoc analysis, with only T1w, MD and E1 having significantly worse performance with respect to the top-performing contrast, SM.

SM and SM1k input data provide the best performance in terms of the mean surface distance (MSD) (Table 5), where full precision floating point values show that SM has a marginally lower MSD value. MK provides the best overall results among the rest of the contrasts, and features a reduced variability across participants. The eigenvalue maps (especially E2 and E3) provide the least informative input contrast to the network in terms of the MSD. MSD shows the same trend shown by the HD95 measure, where the crus accumulates larger errors compared to the rest of the structures. In this case, the statistical significance analysis shows that the differences observed are not significant with respect to SM when measuring MSD values, except for the results obtained when using E1 image input data, which are significantly worse.

**Table 5.**
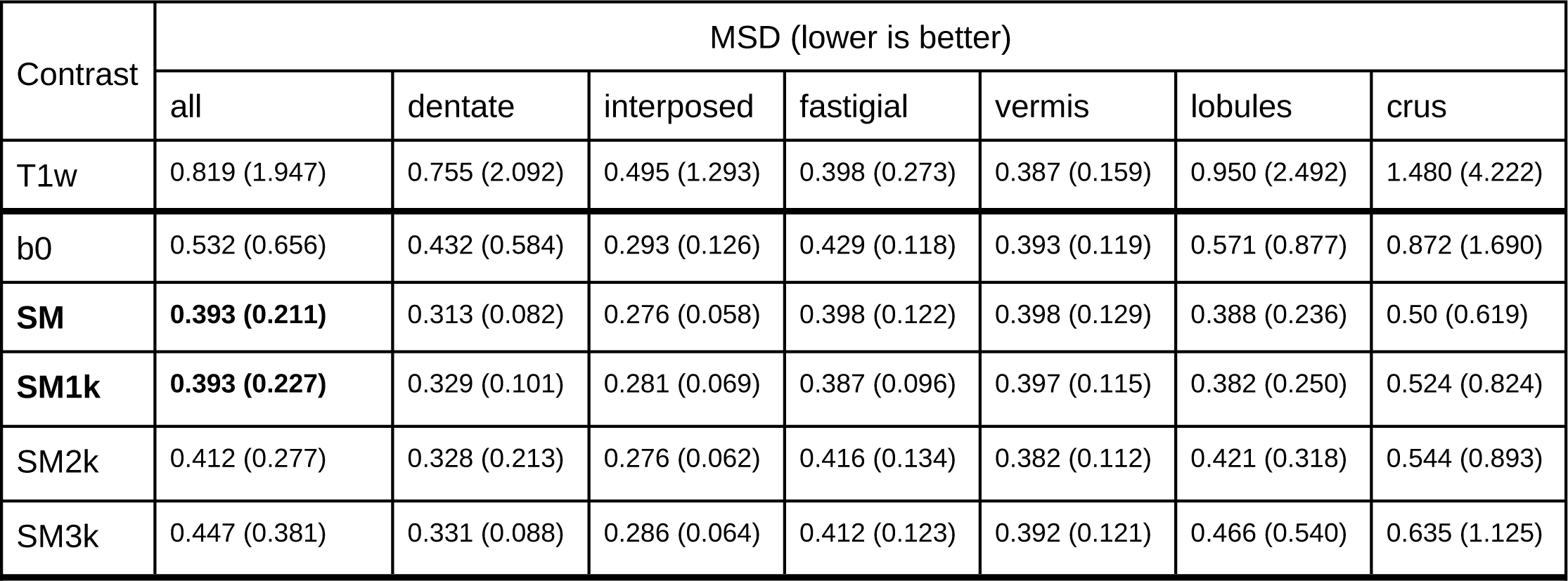
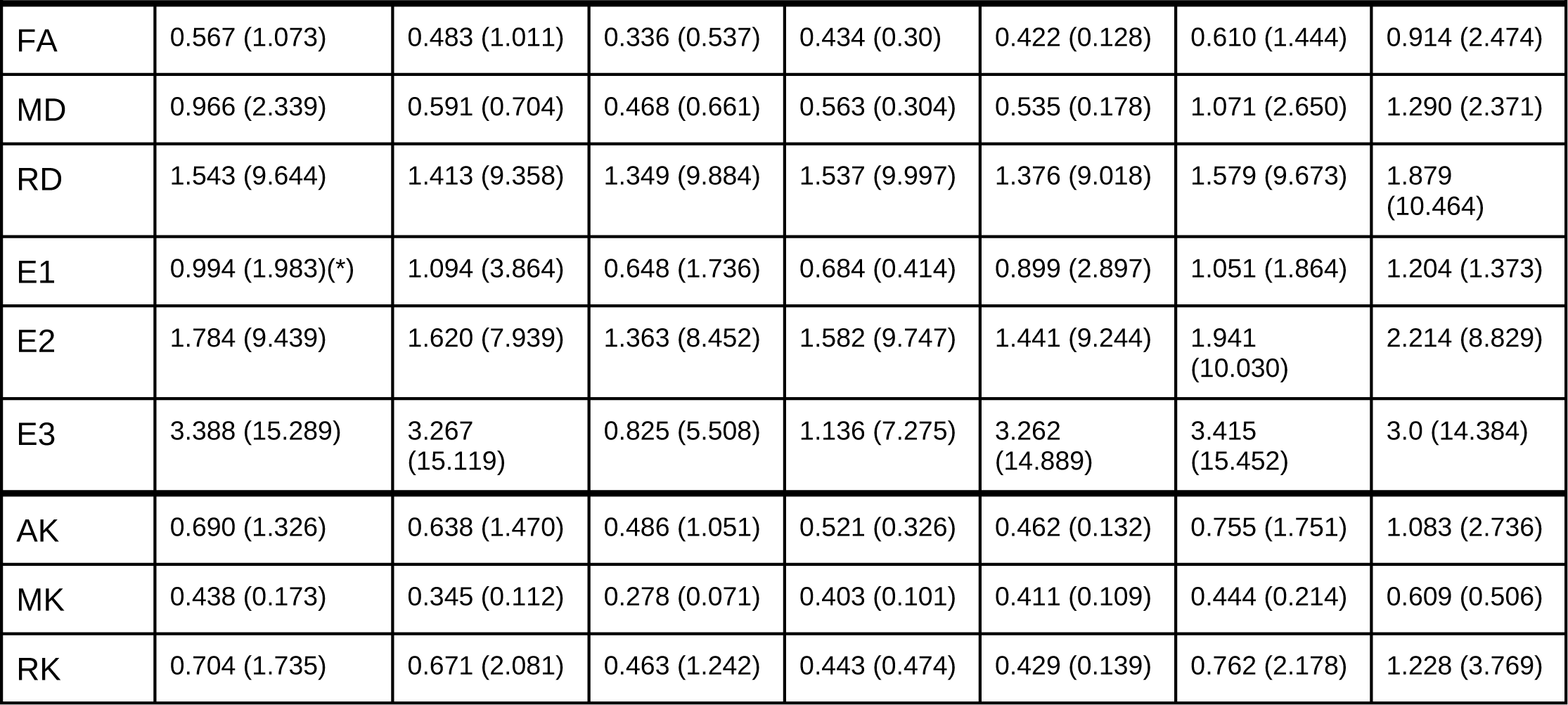
MSD performance across participants. Mean (standard deviation) values across participants for different target structures. Best mean values are highlighted in bold. Asterisks indicate contrasts with significantly worse performance than the top performing contrast.

| Contrast | MSD (lower is better) |  |  |  |  |  |  |
| --- | --- | --- | --- | --- | --- | --- | --- |
|  | all | dentate | interposed | fastigial | vermis | lobules | crus |
| T1w | 0.819 (1.947) | 0.755 (2.092) | 0.495 (1.293) | 0.398 (0.273) | 0.387 (0.159) | 0.950 (2.492) | 1.480 (4.222) |
| b0 | 0.532 (0.656) | 0.432 (0.584) | 0.293 (0.126) | 0.429 (0.118) | 0.393 (0.119) | 0.571 (0.877) | 0.872 (1.690) |
| **SM** | **0.393 (0.211)** | 0.313 (0.082) | 0.276 (0.058) | 0.398 (0.122) | 0.398 (0.129) | 0.388 (0.236) | 0.50 (0.619) |
| **SM1k** | **0.393 (0.227)** | 0.329 (0.101) | 0.281 (0.069) | 0.387 (0.096) | 0.397 (0.115) | 0.382 (0.250) | 0.524 (0.824) |
| SM2k | 0.412 (0.277) | 0.328 (0.213) | 0.276 (0.062) | 0.416 (0.134) | 0.382 (0.112) | 0.421 (0.318) | 0.544 (0.893) |
| SM3k | 0.447 (0.381) | 0.331 (0.088) | 0.286 (0.064) | 0.412 (0.123) | 0.392 (0.121) | 0.466 (0.540) | 0.635 (1.125) |
| FA | 0.567 (1.073) | 0.483 (1.011) | 0.336 (0.537) | 0.434 (0.30) | 0.422 (0.128) | 0.610 (1.444) | 0.914 (2.474) |
| MD | 0.966 (2.339) | 0.591 (0.704) | 0.468 (0.661) | 0.563 (0.304) | 0.535 (0.178) | 1.071 (2.650) | 1.290 (2.371) |
| RD | 1.543 (9.644) | 1.413 (9.358) | 1.349 (9.884) | 1.537 (9.997) | 1.376 (9.018) | 1.579 (9.673) | 1.879 (10.464) |
| E1 | 0.994 (1.983)(*) | 1.094 (3.864) | 0.648 (1.736) | 0.684 (0.414) | 0.899 (2.897) | 1.051 (1.864) | 1.204 (1.373) |
| E2 | 1.784 (9.439) | 1.620 (7.939) | 1.363 (8.452) | 1.582 (9.747) | 1.441 (9.244) | 1.941 (10.030) | 2.214 (8.829) |
| E3 | 3.388 (15.289) | 3.267 (15.119) | 0.825 (5.508) | 1.136 (7.275) | 3.262 (14.889) | 3.415 (15.452) | 3.0 (14.384) |
| AK | 0.690 (1.326) | 0.638 (1.470) | 0.486 (1.051) | 0.521 (0.326) | 0.462 (0.132) | 0.755 (1.751) | 1.083 (2.736) |
| MK | 0.438 (0.173) | 0.345 (0.112) | 0.278 (0.071) | 0.403 (0.101) | 0.411 (0.109) | 0.444 (0.214) | 0.609 (0.506) |
| RK | 0.704 (1.735) | 0.671 (2.081) | 0.463 (1.242) | 0.443 (0.474) | 0.429 (0.139) | 0.762 (2.178) | 1.228 (3.769) |

As shown in Table 6, the center of mass distance (CMD) metric is best when using the various spherical mean contrast data, with SM providing the lowest error, followed by SM1k. MK offers a reasonably low error and stable performance compared to the rest of the contrasts. Eigenvalues and diffusivity-based (MD and RD) contrasts compare unfavorably among the other input data contrasts. CMD follows the trend of the boundary-based measures (HD95, MSD), where the crus structures account for most of the absolute error. From the statistical analysis, it follows that CMD shows minor to non-significant differences when comparing contrasts to the top-performing contrast (SM), with only RD and E3 showing significantly worse performance. This again is analogous to what is shown by the HD95 and MSD boundary-based measure analyses.

**Table 6.** CMD performance across participants. Mean (standard deviation) values across participants for different target structures. Best mean values are highlighted in bold. Asterisks indicate contrasts with significantly worse performance than the top performing contrast.

| Contrast | CMD (lower is better) |  |  |  |  |  |  |
| --- | --- | --- | --- | --- | --- | --- | --- |
|  | all | dentate | interposed | fastigial | vermis | lobules | crus |
| T1w | 1.558 (3.561) | 1.344 (3.560) | 0.864 (1.854) | 0.614 (0.494) | 0.678 (0.275) | 1.832 (4.552) | 2.970 (8.272) |
| b0 | 1.079 (1.501) | 0.775 (1.033) | 0.616 (0.306) | 0.674 (0.237) | 0.742 (0.267) | 1.169 (1.90) | 1.935 (4.562) |
| **SM** | **0.765 (0.478)** | 0.577 (0.240) | 0.608 (0.219) | 0.661 (0.268) | 0.758 (0.284) | 0.764 (0.540) | 1.008 (1.395) |
| SM1k | 0.776 (0.526) | 0.581 (0.222) | 0.586 (0.219) | 0.634 (0.212) | 0.746 (0.257) | 0.782 (0.571) | 1.072 (2.051) |
| SM2k | 0.813 (0.596) | 0.634 (0.360) | 0.605 (0.210) | 0.705 (0.267) | 0.726 (0.248) | 0.842 (0.687) | 1.104 (1.90) |
| SM3k | 0.895 (0.793) | 0.645 (0.251) | 0.629 (0.222) | 0.673 (0.261) | 0.766 (0.279) | 0.939 (1.020) | 1.344 (2.591) |
| FA | 1.140 (2.332) | 0.916 (2.221) | 0.698 (1.073) | 0.691 (0.408) | 0.799 (0.257) | 1.249 (3.082) | 1.954 (5.711) |
| MD | 1.818 (3.769) | 1.129 (1.109) | 0.987 (0.866) | 0.90 (0.431) | 1.073 (0.437) | 1.981 (3.923) | 2.578 (4.856) |
| RD | 2.272 (11.093)(*) | 2.001 (10.347) | 1.819 (10.291) | 1.861 (9.912) | 1.926 (9.905) | 2.378 (11.495) | 3.024 (12.733) |
| E1 | 1.863 (2.991) | 1.809 (4.637) | 1.317 (2.230) | 1.084 (0.601) | 1.637 (3.914) | 2.021 (3.172) | 2.320 (2.674) |
| E2 | 2.767 (10.903) | 2.497 (9.366) | 2.038 (9.307) | 1.868 (9.767) | 2.009 (10.096) | 3.032 (11.532) | 4.060 (12.741) |
| E3 | 3.859 (15.686)<br>(**) | 3.659 (15.386) | 1.171 (5.758) | 1.429 (7.524) | 3.695 (15.296) | 3.904 (15.917) | 3.538 (14.249) |
| AK | 1.442 (2.992) | 1.265 (3.079) | 0.984 (1.537) | 0.879 (0.505) | 0.938 (0.318) | 1.60 (3.877) | 2.375 (7.092) |
| MK | 0.836 (0.379) | 0.634 (0.262) | 0.574 (0.207) | 0.659 (0.215) | 0.780 (0.261) | 0.855 (0.467) | 1.198 (1.209) |
| RK | 1.395 (3.361) | 1.220 (3.460) | 0.859 (1.644) | 0.718 (0.619) | 0.824 (0.291) | 1.531 (4.208) | 2.543 (8.131) |

SM1k provides the best performance in terms of the volume similarity (VS) (Table 7). The eigenvalue images (E1, E2, E3), b0, T1w and MD data provide the least competitive performance for VS. When comparing results against the top-performing contrast (SM1k), the post hoc statistical analysis reveals that all contrasts perform significantly worse, with the exception of SM.

**Table 7.** VS performance across participants. Mean (standard deviation) values across participants for different target structures. Best mean values are highlighted in bold. Asterisks indicate contrasts with significantly worse performance than the top performing contrast.

| Contrast | VS (higher is better) |  |  |  |  |  |  |
| --- | --- | --- | --- | --- | --- | --- | --- |
|  | all | dentate | interposed | fastigial | vermis | lobules | crus |
| T1w | 0.937 (0.062)(***) | 0.952 (0.083) | 0.931 (0.074) | 0.702 (0.232) | 0.945 (0.022) | 0.956 (0.079) | 0.953 (0.111) |
| b0 | 0.940 (0.042)(***) | 0.947 (0.056) | 0.942 (0.041) | 0.691 (0.227) | 0.941 (0.022) | 0.964 (0.051) | 0.968 (0.070) |
| SM | 0.958 (0.017) | 0.970 (0.019) | 0.950 (0.033) | 0.812 (0.183) | 0.948 (0.018) | 0.975 (0.018) | 0.979 (0.030) |
| **SM1k** | **0.961 (0.018)** | 0.953 (0.042) | 0.941 (0.041) | 0.895 (0.056) | 0.946 (0.022) | 0.976 (0.018) | 0.978 (0.051) |
| SM2k | 0.952 (0.022)(**) | 0.965 (0.027) | 0.950 (0.030) | 0.720 (0.230) | 0.948 (0.025) | 0.975 (0.017) | 0.976 (0.036) |
| SM3k | 0.954 (0.023)(*) | 0.963 (0.029) | 0.946 (0.034) | 0.797 (0.190) | 0.947 (0.019) | 0.972 (0.027) | 0.973 (0.051) |
| FA | 0.950 (0.040)(*) | 0.966 (0.021) | 0.948 (0.039) | 0.876 (0.079) | 0.922 (0.051) | 0.967 (0.057) | 0.970 (0.068) |
| MD | 0.928 (0.097)(**) | 0.941 (0.114) | 0.915 (0.122) | 0.762 (0.216) | 0.923 (0.098) | 0.945 (0.102) | 0.951 (0.092) |
| RD | 0.948 (0.060)(*) | 0.958 (0.041) | 0.924 (0.084) | 0.870 (0.132) | 0.936 (0.064) | 0.960 (0.059) | 0.967 (0.050) |
| E1 | 0.930 (0.064)(***) | 0.924 (0.110) | 0.902 (0.092) | 0.828 (0.151) | 0.920 (0.051) | 0.944 (0.075) | 0.960 (0.043) |
| E2 | 0.925 (0.078)(***) | 0.919 (0.113) | 0.912 (0.097) | 0.748 (0.243) | 0.930 (0.060) | 0.941 (0.085) | 0.950 (0.076) |
| E3 | 0.919 (0.181)(*) | 0.929 (0.184) | 0.927 (0.177) | 0.768 (0.229) | 0.912 (0.174) | 0.936 (0.189) | 0.939 (0.194) |
| AK | 0.951 (0.024)(**) | 0.958 (0.034) | 0.925 (0.047) | 0.873 (0.087) | 0.937 (0.022) | 0.965 (0.027) | 0.970 (0.037) |
| MK | 0.956 (0.014)(*) | 0.959 (0.024) | 0.929 (0.041) | 0.885 (0.060) | 0.941 (0.020) | 0.971 (0.015) | 0.975 (0.030) |
| RK | 0.945 (0.037)(***) | 0.964 (0.027) | 0.945 (0.032) | 0.706 (0.233) | 0.946 (0.019) | 0.967 (0.044) | 0.966 (0.076) |

The label detection rate (LDR) metric exhibits the best results for SM1k and the DKI-derived AK and MK input data (Table 8). The lowest detection rates are recorded for the fastigial nuclei, as they are notably small structures at the considered spatial image resolution.

**Table 8.** LDR performance across participants. Mean (standard deviation) values across participants for different target structures. Best mean values are highlighted in bold.

| Contrast | LDR (higher is better) |  |  |  |  |  |  |
| --- | --- | --- | --- | --- | --- | --- | --- |
|  | all | dentate | interposed | fastigial | vermis | lobules | crus |
| T1w | 0.988 (0.015) | 1.0 (0.0) | 0.995 (0.05) | 0.795 (0.247) | 1.0 (0.0) | 1.0 (0.0) | 1.0 (0.0) |
| b0 | 0.988 (0.015) | 1.0 (0.0) | 1.0 (0.0) | 0.8 (0.246) | 1.0 (0.0) | 0.999 (0.006) | 1.0 (0.0) |
| SM | 0.994 (0.012) | 1.0 (0.0) | 1.0 (0.0) | 0.9 (0.201) | 1.0 (0.0) | 1.0 (0.0) | 1.0 (0.0) |
| **SM1k** | **1.0 (0.003)** | 1.0 (0.0) | 1.0 (0.0) | 1.0 (0.0) | 0.999 (0.012) | 1.0 (0.0) | 1.0 (0.0) |
| SM2k | 0.989 (0.014) | 1.0 (0.0) | 1.0 (0.0) | 0.8 (0.246) | 1.0 (0.0) | 1.0 (0.0) | 1.0 (0.0) |
| SM3k | 0.994 (0.012) | 1.0 (0.0) | 1.0 (0.0) | 0.9 (0.201) | 1.0 (0.0) | 1.0 (0.0) | 1.0 (0.0) |
| FA | 0.994 (0.012) | 1.0 (0.0) | 1.0 (0.0) | 1.0 (0.0) | 0.975 (0.050) | 1.0 (0.006) | 1.0 (0.0) |
| MD | 0.987 (0.068) | 0.99 (0.1) | 0.99 (0.01) | 0.89 (0.220) | 0.989 (0.101) | 0.994 (0.057) | 1.0 (0.0) |
| RD | 0.999 (0.012) | 1.0 (0.0) | 1.0 (0.0) | 0.99 (0.070) | 0.998 (0.025) | 0.999 (0.006) | 1.0 (0.0) |
| E1 | 0.998 (0.018) | 0.995 (0.05) | 0.995 (0.05) | 0.99 (0.1) | 0.999 (0.013) | 0.999 (0.009) | 1.0 (0.0) |
| E2 | 0.993 (0.013) | 1.0 (0.0) | 1.0 (0.0) | 0.89 (0.208) | 0.999 (0.013) | 1.0 (0.0) | 1.0 (0.0) |
| E3 | 0.976 (0.098) | 0.995 (0.05) | 0.97 (0.171) | 0.87 (0.252) | 0.986 (0.069) | 0.984 (0.097) | 0.97 (0.152) |
| AK | **1.0 (0.003)** | 1.0 (0.0) | 1.0 (0.0) | 0.995 (0.05) | 1.0 (0.0) | 1.0 (0.0) | 1.0 (0.0) |
| MK | **1.0 (0.0)** | 1.0 (0.0) | 1.0 (0.0) | 1.0 (0.0) | 1.0 (0.0) | 1.0 (0.0) | 1.0 (0.0) |
| RK | 0.988 (0.015) | 1.0 (0.0) | 1.0 (0.0) | 0.795 (0.247) | 1.0 (0.0) | 1.0 (0.0) | 1.0 (0.0) |

#### 3.1.2. Overall performance

For each metric, contrasts were ranked according to their mean values across all labels and participants. These rank values were then averaged to provide an overall ranking across metrics, shown in Table 9. Rankings range from best to worst (1 to 15). Results show that the b=1000 s/mm^2^ shell spherical mean (SM1k) data rank best, with the rest of the spherical mean contrasts (i.e., SM, SM2k, SM3k) being in the top five. The DKI-derived mean kurtosis (MK) ranks third, following SM1k and SM.

**Table 9.**
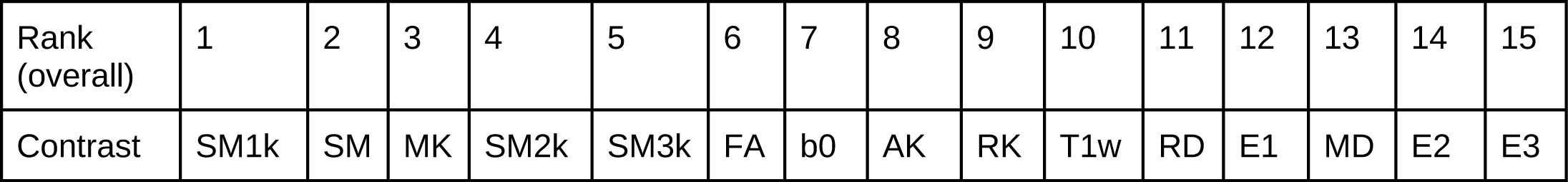
Overall performance ranking across metrics. Best (1) to worst (15).

|  |  |  |  |  |  |  |  |  |  |  |  |  |  |  |  |
| --- | --- | --- | --- | --- | --- | --- | --- | --- | --- | --- | --- | --- | --- | --- | --- |
| Rank (overall) | 1 | 2 | 3 | 4 | 5 | 6 | 7 | 8 | 9 | 10 | 11 | 12 | 13 | 14 | 15 |
| Contrast | SM1k | SM | MK | SM2k | SM3k | FA | b0 | AK | RK | T1w | RD | E1 | MD | E2 | E3 |

#### 3.1.3. Qualitative analysis

##### 3.1.3.1. Visualization of predicted segmentations

Figure 3 shows coronal anterior views of the predicted segmentation labels for all considered contrasts on a given participant. Sagittal and axial views of the predicted labels for each contrast are shown in the Appendix section.

**Figure 3.**
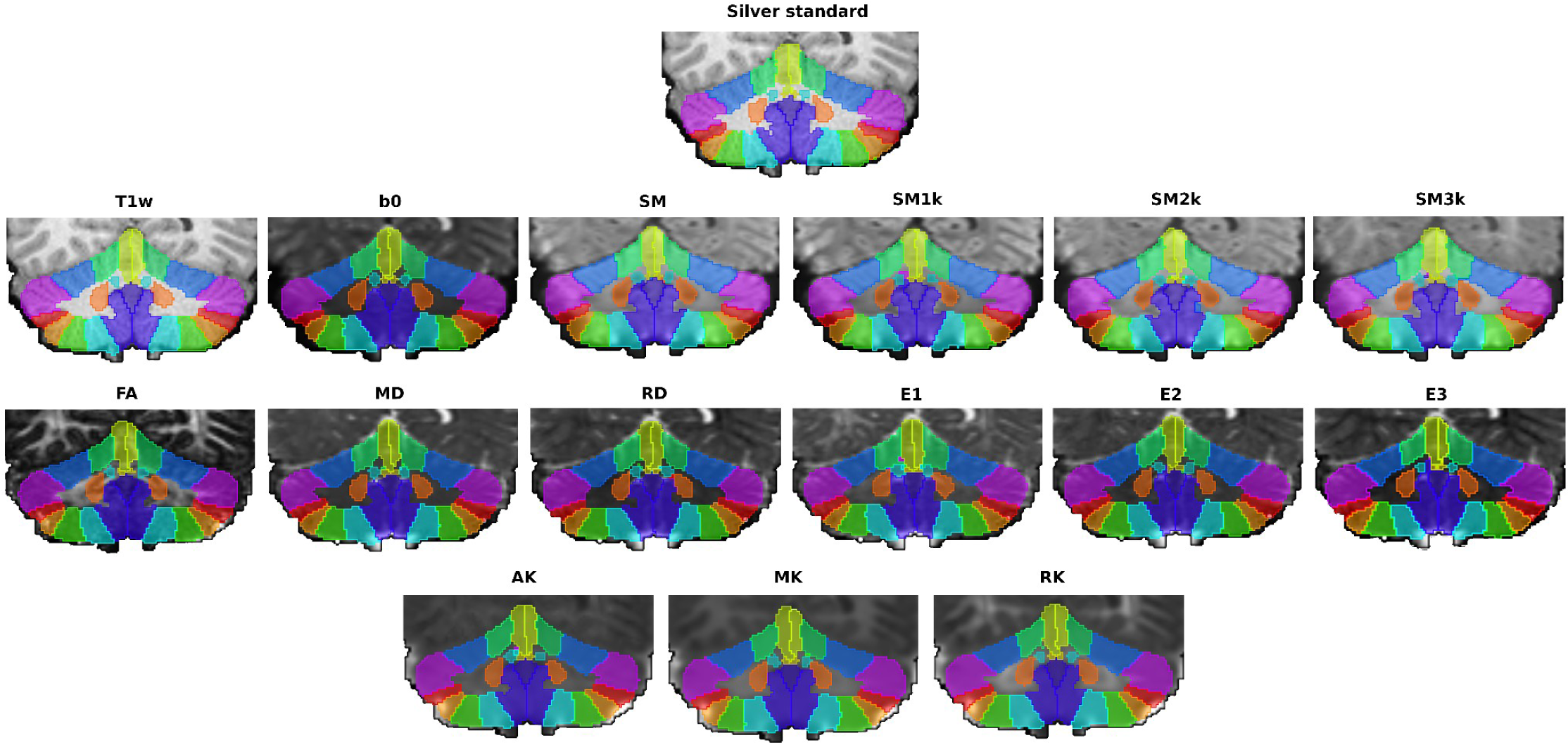
Predicted segmentations for a given participant. All coronal anterior views.

##### 3.1.3.2. Detailed visualization of top-performing contrasts

Following the quantitative results, we illustrate the differences between the silver standard and the predicted segmentations using the top-performing contrast images, the multi-shell (SM) and b=1000 s/mm^2^ (SM1k) spherical means. In addition to the predictions on the SM and SM1k input data, we show the predictions obtained from the T1w images for reference, as T1w data are commonly employed by popular segmentation tools. The following figures provide visualizations of segmentations of the deep cerebellar nuclei and of the cerebellar cortical regions, i.e. the lobules, vermis and crus.

Figure 4 provides volumetric illustrations of the silver standard deep cerebellar nuclei and the predictions obtained using the T1w, SM and SM1k data, respectively, for a given participant. As shown, in this example participant both predictions on SM and SM1k data are able to recover all nuclei, but predicting on T1w misses the left fastigial nucleus.

**Figure 4.**
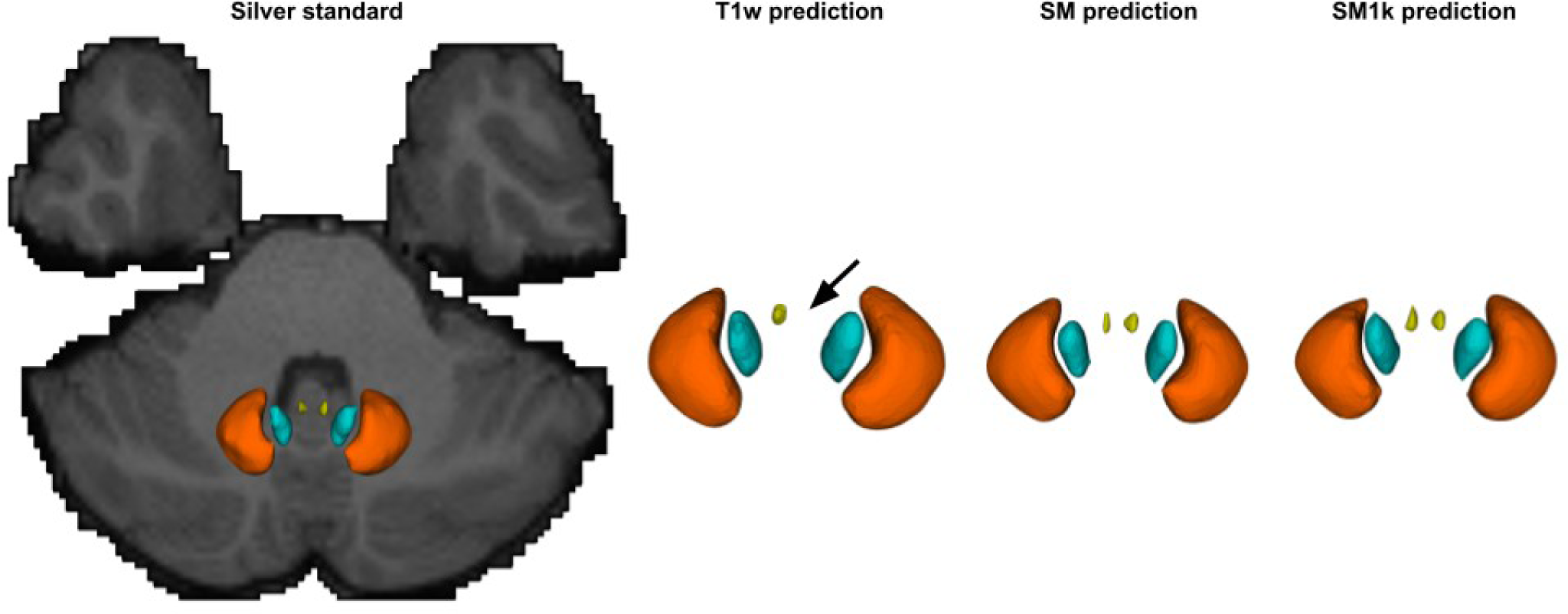
Silver standard deep cerebellar nuclei and the predictions obtained using the T1w, SM and SM1k data on a given participant. The black arrow points at the left fastigial nucleus, which is missed by the prediction on T1w data. Colors and nuclei: orange: dentate; emerald: interposed; lime: fastigial. All axial superior views.

Figure 5 shows contour plots of the left and right dentate nuclei for two participants for the T1w, SM and SM1k contrasts. Qualitatively and overall, both the SM and SM1k contrasts follow more closely the contour of the silver standard.

**Figure 5.**
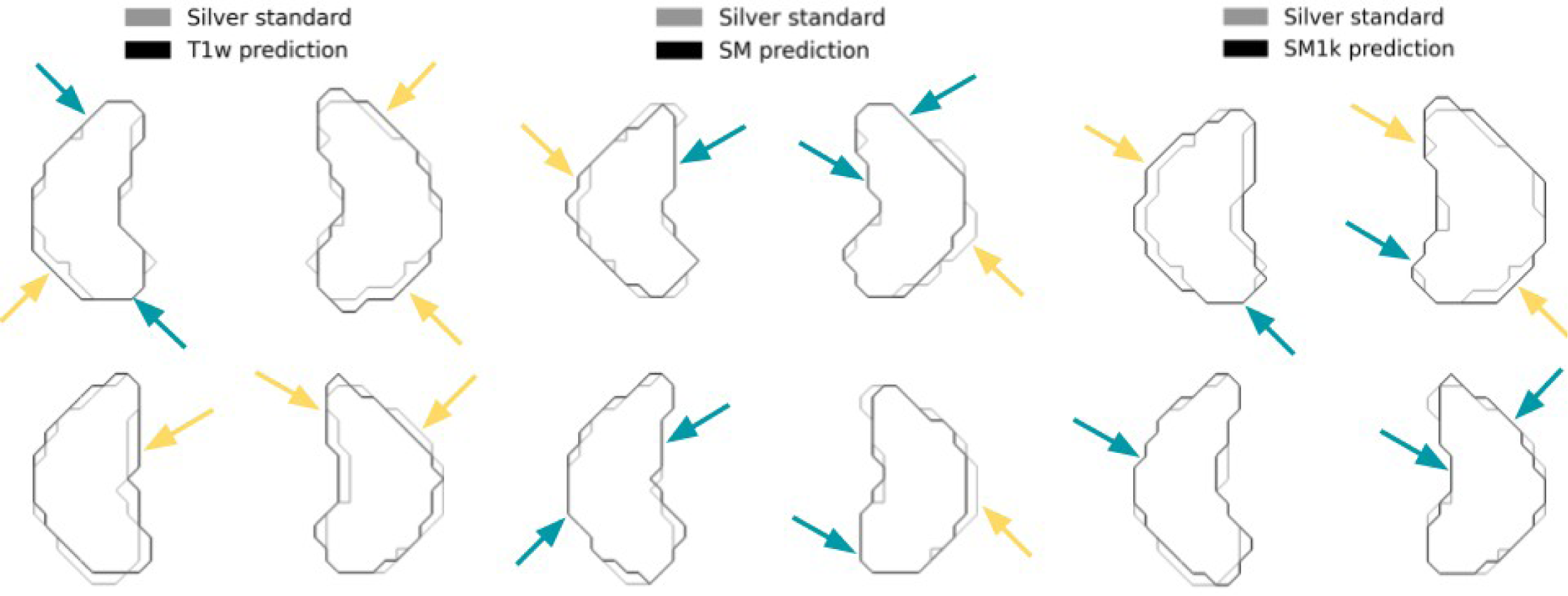
Axial view contour plots of the left and right dentate nuclei: silver standard and predicted contours for two test participants (upper row; lower row) for the T1w, SM and SM1k contrasts (left to right). Emerald arrows point to regions where the prediction matches more closely the silver standard; yellow arrows point to regions where the prediction shows a larger deviation from the silver standard.

Figure 6 shows flat surface maps of the cerebellar cortical regions on two test participants as predicted on T1w, SM and SM1k data, respectively. Although all three predictions seem to provide a reasonable agreement with the silver standard, some subtle differences (indicated with black arrows) are apparent in certain regions across the predictions.

**Figure 6.**
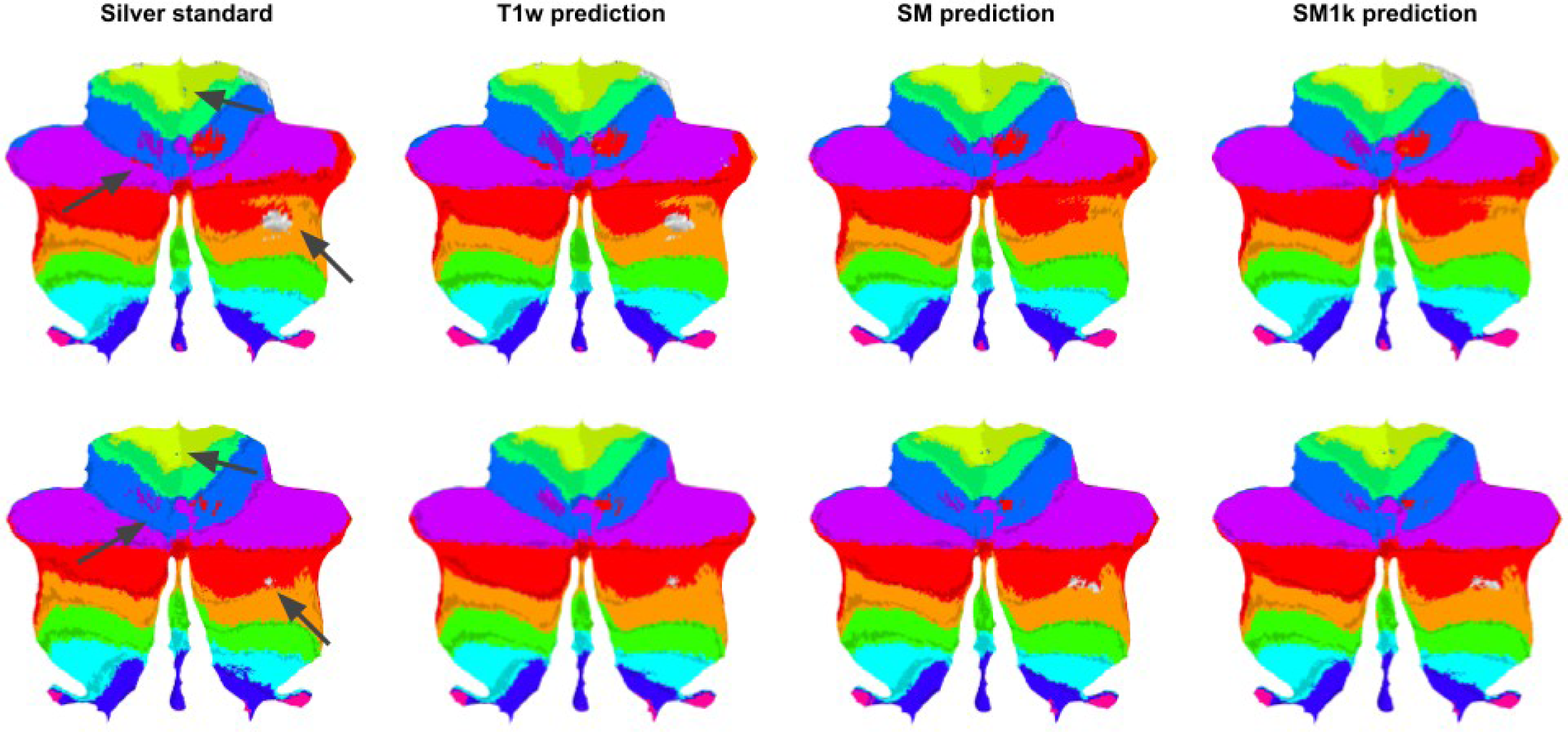
Surface flat maps of the cerebellar regions for two test participants (upper row; lower row) for the silver standard, and the predicted label maps using the T1w, SM and SM1k contrasts (left to right). Dark arrows point to regions where predictions show the largest differences across contrasts.

### 3.2. Experiment 2: Generalization of models to fewer gradient directions

The generalization ability of the model is analyzed by testing the models on the SM and SM1k contrasts computed from data with lower numbers of diffusion encoding gradient directions (20, 30 and 60), as described in section 2.5.2. In this experiment, we focus on the top-performing contrasts, SM and SM1k, following our overall ranking results (Table 9). The performance metrics of the predicted segmentations (across all labels) on the subsampled data are shown in Table 10. From these results, it can be concluded that the performance is stable within the range of gradient directions that is typical in dMRI, including the lower-end, clinical-style 20 directions. Compared to the results using all directions, the performances are comparable, with only slightly worse performance in MSD (0.4 vs. 0.39 on both SM and SM1k) and CMD (0.78 vs 0.77 and 0.8 vs 0.78 for SM and SM1k, respectively).

**Table 10.** Predicted segmentation performance for spherical mean input data computed on DWI multi-shell and b=1000 s/mm2 data subsampled at 20, 30 and 60 directions. “all” values are repeated here from the corresponding previous tables to serve as a reference.

| Contrast | Directions | DSC | HD95 | MSD | CMD | VS | LDR |
| --- | --- | --- | --- | --- | --- | --- | --- |
| SM | 20 | 0.837 (0.032) | 1.462 (0.932) | 0.40 (0.213) | 0.782 (0.483) | 0.957 (0.017) | 0.994 (0.012) |
|  | 30 | 0.838 (0.032) | 1.448 (0.929) | 0.399 (0.213) | 0.779 (0.485) | 0.957 (0.017) | 0.994 (0.012) |
|  | 60 | 0.838 (0.032) | 1.461 (0.932) | 0.399 (0.211) | 0.779 (0.479) | 0.957 (0.017) | 0.994 (0.012) |
|  | all | 0.840 (0.032) | 1.443 (0.911) | 0.393 (0.211) | 0.765 (0.478) | 0.958 (0.017) | 0.994 (0.012) |
| SM1k | 20 | 0.843 (0.031) | 1.460 (1.045) | 0.401 (0.258) | 0.797 (0.622) | 0.960 (0.021) | 1.0 (0.003) |
|  | 30 | 0.843 (0.031) | 1.459 (1.054) | 0.401 (0.263) | 0.797 (0.637) | 0.960 (0.021) | 1.0 (0.003) |
|  | 60 | 0.843 (0.031) | 1.459 (1.060) | 0.402 (0.266) | 0.798 (0.637) | 0.960 (0.021) | 1.0 (0.003) |
|  | all | 0.844 (0.030) | 1.446 (0.979) | 0.393 (0.227) | 0.776 (0.526) | 0.961 (0.018) | 1.0 (0.003) |

## 4. Discussion

The fine-grained segmentation of brain tissue structures is an essential step towards supplying increasingly accurate anatomically informed analyses, including, for example, white matter dMRI tractography. Most often, structural acquisitions (T1w or T2w) are used to segment the brain tissues. Such segmentation maps are then registered to dMRI acquisitions in order to proceed with diffusion-specific analyses. However, segmentation errors and registration misalignments have an impact on the accuracy of results. Thus, it is desirable to obtain the best possible segmentations on the native dMRI images in order to reduce the error sources. In this work we compare the performance of different dMRI-derived contrasts and evaluate them on the cerebellar segmentation task using a neural network model. We specifically choose to use contrasts derived from both the diffusion signal and local parameterizations (DTI and DKI).

We show in our experiments that the spherical mean contrasts obtained as the averaged diffusion-weighted signal perform particularly well compared to other dMRI-derived contrasts that have been used extensively in the literature to train DL models. The spherical mean contrast is easy to compute and can be used in single- or multi-shell acquisition schemes. Experiments show that spherical mean contrasts provide stable performance across overlap-, boundary-, and property-based metrics: the multi-shell and b=1000 s/mm^2^ shell spherical means (SM and SM1k, respectively) rank consistently as the top performers. A statistical analysis reveals that for the overlap- and property-based metrics, spherical mean images provide significantly superior performance. The boundary-based metrics present larger standard deviation values across participants, and thus may explain the general lack of significant differences in performance across methods as shown by these metrics. The metrics computed on the segmentations obtained using spherical mean data also exhibit lower variance across participants compared to the variance of the rest of the contrasts. As demonstrated by the LDR (label detection rate) metric (Table 8) and as shown in Figure 4, the spherical mean contrasts reliably detect all labels, despite some of these presenting a challenging task due to their reduced size (particularly the fastigial nuclei, which are the smallest nuclei structures studied in this work). This may be attributed to the relatively high image contrast between the darker DCNs (e.g., dentate) and the surrounding white matter, as shown in Figure 2. This differs for example from the rather homogeneous intensity seen around the DCNs on the T1w contrast images, which may entail increased segmentation difficulty for the neural network or for human experts. Given the appropriate spatial resolution and/or protocol, spherical mean data may thus enable manual contouring of deep cerebellar structures in order to compare automated methods to expert-delineated segmentations.

Regarding the other contrasts studied, the spherical means computed over the b=2000 s/mm^2^ and b=3000 s/mm^2^ shell data (SM2k and SM3k) also compare favorably to most other dMRI-derived contrasts, but underperform SM and SM1k. As the diffusion weighting (b-value) increases, the signal decays significantly, and thus the SNR of the acquired signal decreases (Jones & Basser, 2004; Magin, 2016), which may explain the performance decline using SM2k and SM3k data. Some of the studied contrasts exhibit a large variability in performance across participants: in particular, when using the eigenvalue maps as the input contrast, the model is not able to segment the target structures on some participants.

A gradient weighting of b=1000 s/mm^2^ is a well-established value for several clinical applications (Descoteaux, 2015; Hagmann et al., 2006), and thus, segmentation of cerebellar structures on retrospective clinical data may be facilitated using spherical mean contrast (SM1k) input to a neural network. Different authors (Jones, 2004; Tournier et al., 2013) have studied the lower bounds for gradient sampling directions for meaningful local orientation property detection using different models. Here we set 20 directions as a practical lower bound for DTI-like acquisitions. The present experiments show that the spherical mean contrasts provide good results in low angular resolution settings (Table 10): the segmentation performance is stable across a range of diffusion-encoding gradient directions that optimally cover the angular space. This suggests that the cerebellar target structures may be segmented successfully when scan time is limited (e.g. clinical settings), and when using as few as 20 diffusion-encoding gradient directions.

### 4.1. Limitations and future work

Here we discuss limitations of the current work and potential avenues for further research to leverage the use of spherical mean data in cerebellar tissue segmentation and analysis. First, in this work, we perform a comprehensive evaluation of individual dMRI-derived image contrasts for cerebellar segmentation. Future work may explore the usage of multiple input image contrasts, which may provide a more accurate delineation of gray and white matter structures (e.g. (F. Zhang et al., 2021, 2024)). Future work may also explore the usage of tractography-derived contrasts, such as track density imaging, which can help in delineation of brain structures such as the thalamic nuclei (Basile et al., 2021; Semedo et al., 2018). Second, future work may explore various strategies to potentially improve model performance, such as test-time augmentation (G. Wang et al., 2019) or multi-resolution segmentation strategies (Morell-Ortega et al., 2024). Third, in this work we rely on the labels provided by a single tool as the silver standard to train our models. A growing body of work exists in the deep learning domain that considers including some form of uncertainty-based measure to improve the model outcomes (see, for example, (Ji et al., 2021; L. Zhang et al., 2023)). Such strategies comprise probabilistic approaches to model the variability from multiple annotations (also referred to as “noisy labels”) of the same target to produce either a single, consensus segmentation or multiple outputs for the target. Introducing such enhancements to the dMRI-derived contrast segmentation framework is left for future study. Similarly, further work is required to investigate the impact of the proposed segmentation strategy in downstream tasks, such as dMRI tractography. An improved segmentation outcome could allow, for example, the application of more fine-grained anatomical constraints to tractography methods, which should provide more accurate connectivity results. Finally, in a future work, model performance using spherical mean contrast data could be compared to segmentations obtained from SWI and QSM data in order to assess results against acquisitions genuinely targeted towards capturing biological contrast in the deep cerebellar gray matter nuclei.

## 5. Conclusion

In this work, we investigate dMRI-derived contrasts for cerebellar tissue segmentation using a deep learning model. We provide evidence that the spherical mean contrast provides significantly improved results over other dMRI-derived contrasts that are used in brain tissue segmentation. Improved dMRI-based segmentation can be useful to overcome limitations from structural data segmentations, including inaccuracies introduced by registration processes. Thus, we believe that future work focusing on using dMRI data for brain structure segmentation purposes would benefit from including the spherical mean in their experiments.

## Acknowledgements

We gratefully acknowledge funding provided by the following grants: National Institutes of Health (NIH) grants R01MH132610, R01MH125860, R01MH119222, R01NS125307, R01NS125781. FZ is in part supported by National Key R&D Program of China (No. 2023YFE0118600), and the National Natural Science Foundation of China (No. 62371107)

Data were provided (in part) by the Human Connectome Project, WU-Minn Consortium (Principal Investigators: David Van Essen and Kamil Ugurbil; 1U54MH091657) funded by the 16 NIH Institutes and Centers that support the NIH Blueprint for Neuroscience Research; and by the McDonnell Center for Systems Neuroscience at Washington University. We thank the Enterprise Research Infrastructure & Services at Mass General Brigham for their support and for the provision of the Scientific Computing (SciC) Linux Clusters. This work used the Jetstream2 on-demand computing infrastructure (Hancock et al., 2021) at Indiana University through allocation MED230035 from the Advanced Cyberinfrastructure Coordination Ecosystem: Services & Support (ACCESS) program, which is supported by National Science Foundation grants #2138259, #2138286, #2138307, #2137603, and #2138296 (Boerner et al., 2023). The code used for the experiments will be made available on https://github.com/SlicerDMRI upon acceptance of the manuscript.

## A. Appendix

### A.1. Neural network details

Figure 7 shows the neural network architecture used in this work. The network follows the SegResNet backbone proposed in (Myronenko, 2019). Following evidence (Woo & Lee, 2021) about the benefits of using fully volumetric (3D) spatial contexts for brain tissue segmentation, we choose to accept whole-brain 3D images at the input of the network. In order to be able to attribute the performance differences to the input contrast, we choose to accept full-sized 3D images (192x192x192) at the input of our network. Following this choice, we do not regularize the model in order to fit our hardware budget constraints. The network accepts 3D images and employs 1, 2 (twice) and 4 downsampling blocks in the contracting layers, and a single upsampling block in the expanding layers.

**Figure 7.**
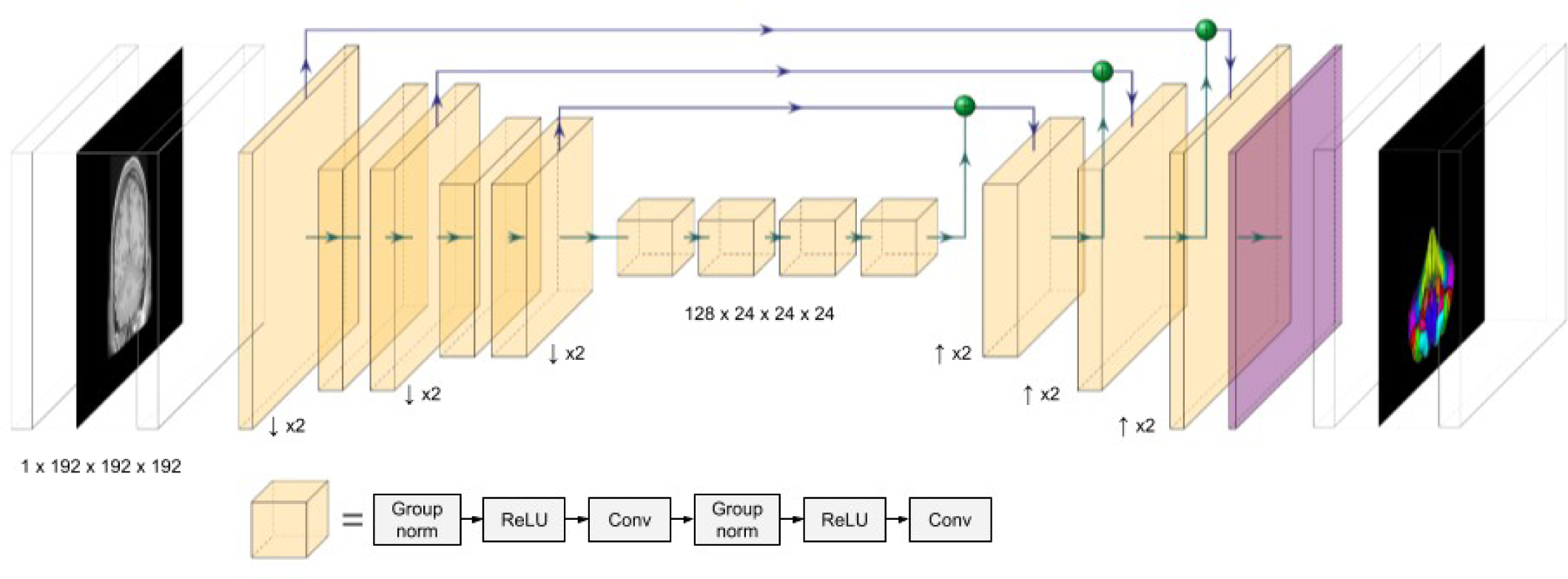
Neural network architecture used in this work.

### A.2. Cerebellar segmentation structures

The silver standard in this work considers the following cerebellar structures: Vermis_VI; Vermis_CrusI; Vermis_CrusII; Vermis_VIIb; Vermis_VIIIa; Vermis_VIIIb; Vermis_IX; Vermis_X; Left_I_IV; Left_V; Left_VI; Left_CrusI; Left_CrusII; Left_VIIb; Left_VIIIa; Left_VIIIb; Left_IX; Left_X; Left_Dentate; Left_Interposed; Left_Fastigial; Right_I_IV; Right_V; Right_VI; Right_CrusI; Right_CrusII; Right_VIIb; Right_VIIIa; Right_VIIIb; Right_IX; Right_X; Right_Dentate; Right_Interposed; Right_Fastigial.

### A.3. Additional segmentation views

Sagittal left (Figure 8) and axial superior (Figure 9) views of the predicted segmentation labels for all considered contrasts on the same participant shown in Figure 3.

**Figure 8.**
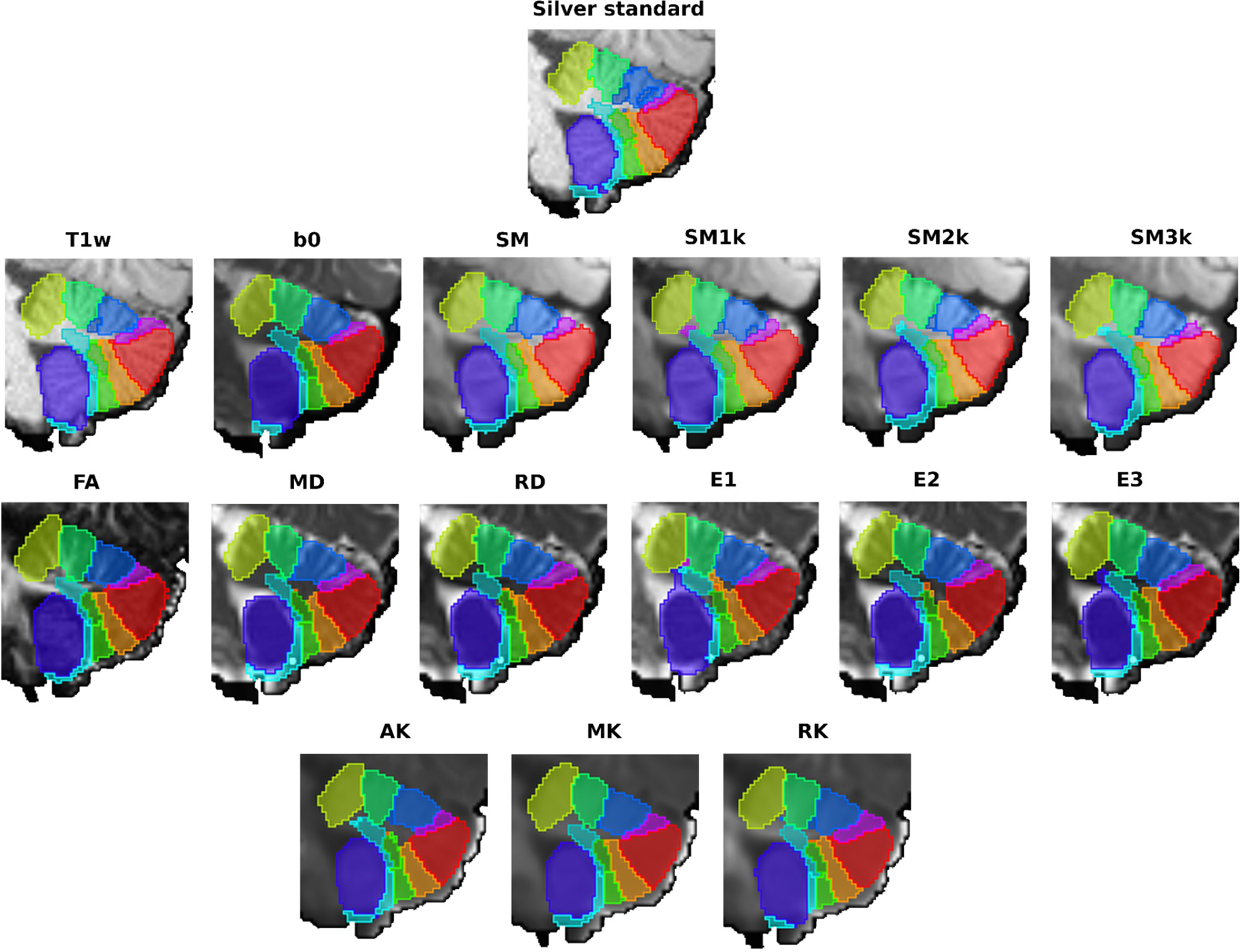
Predicted segmentations for a given participant. All sagittal left views.

**Figure 9.**
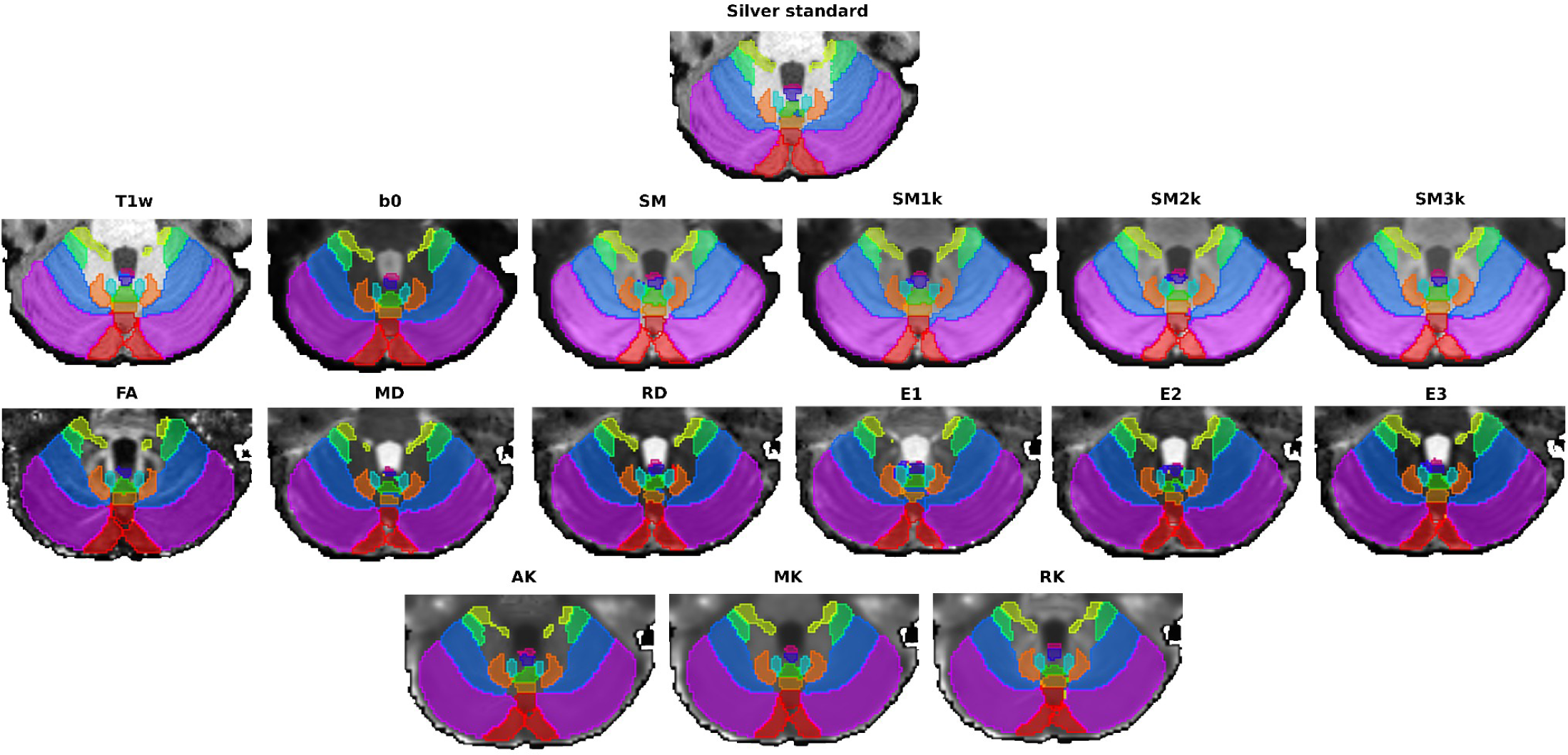
Predicted segmentations for a given participant. All axial superior views.

### Footnotes

1 dMRI and DWI are used interchangeably in literature to refer to diffusion(-weighted) MRI. In the particular context of this work we use DWI to refer to the diffusion(-weighted) data volumes, i.e. as issued by any denoising or distortion correction pre-processing pipeline.

2 In the absence of ground truth annotations, the term “silver standard” is used to refer to annotations that act as a valid substitute for specific purposes. In the context of this work, we consider the segmentations provided by SUIT to be our “silver standard”.

3 The BraTS 2021 Challenge leaderboard is available at https://www.synapse.org/Synapse:syn25829067/wiki/611504

## References

Altman, J., & Bayer, S. A. (1996). Development of the cerebellar system. CRC Press.

Azevedo, F. A. C., Carvalho, L. R. B., Grinberg, L. T., Farfel, J. M., Ferretti, R. E. L., Leite, R. E. P., Jacob Filho, W., Lent, R., & Herculano-Houzel, S. (2009). Equal numbers of neuronal and nonneuronal cells make the human brain an isometrically scaled-up primate brain. The Journal of Comparative Neurology, 513(5), 532–541.

Basile, G. A., Bertino, S., Bramanti, A., Ciurleo, R., Anastasi, G. P., Milardi, D., & Cacciola, A. (2021). In Vivo Super-Resolution Track-Density Imaging for Thalamic Nuclei Identification. Cerebral Cortex, 31(12), 5613–5636.

Baumann, O., Borra, R. J., Bower, J. M., Cullen, K. E., Habas, C., Ivry, R. B., Leggio, M., Mattingley, J. B., Molinari, M., Moulton, E. A., Paulin, M. G., Pavlova, M. A., Schmahmann, J. D., & Sokolov, A. A. (2015). Consensus paper: the role of the cerebellum in perceptual processes. Cerebellum, 14(2), 197–220.

Beliveau, V., Nørgaard, M., Birkl, C., Seppi, K., & Scherfler, C. (2021). Automated segmentation of deep brain nuclei using convolutional neural networks and susceptibility weighted imaging. Human Brain Mapping, 42(15), 4809–4822.

Boerner, T. J., Deems, S., Furlani, T. R., Knuth, S. L., & Towns, J. (2023). ACCESS: Advancing Innovation: NSF’s Advanced Cyberinfrastructure Coordination Ecosystem: Services & Support. Practice and Experience in Advanced Research Computing, 173–176.

Buckner, R. L. (2013). The cerebellum and cognitive function: 25 years of insight from anatomy and neuroimaging. Neuron, 80(3), 807–815.

Carass, A., Cuzzocreo, J. L., Han, S., Hernandez-Castillo, C. R., Rasser, P. E., Ganz, M., Beliveau, V., Dolz, J., Ben Ayed, I., Desrosiers, C., Thyreau, B., Romero, J. E., Coupé, P., Manjón, J. V., Fonov, V. S., Collins, D. L., Ying, S. H., Onyike, C. U., Crocetti, D., … Prince, J. L. (2018). Comparing fully automated state-of-the-art cerebellum parcellation from magnetic resonance images. NeuroImage, 183, 150–172.

Caruyer, E., Lenglet, C., Sapiro, G., & Deriche, R. (2013). Design of multishell sampling schemes with uniform coverage in diffusion MRI. Magnetic Resonance in Medicine: Official Journal of the Society of Magnetic Resonance in Medicine / Society of Magnetic Resonance in Medicine, 69(6), 1534–1540.

Cheng, H., Newman, S., Afzali, M., Fadnavis, S. S., & Garyfallidis, E. (2020). Segmentation of the brain using direction-averaged signal of DWI images. Magnetic Resonance Imaging, 69, 1–7.

Ciritsis, A., Boss, A., & Rossi, C. (2018). Automated pixel-wise brain tissue segmentation of diffusion-weighted images via machine learning. NMR in Biomedicine, 31(7), e3931.

Creators MONAI Consortium. (n.d.). MONAI: Medical Open Network for AI. 10.5281/zenodo.8436376

Descoteaux, M. (2015). High angular resolution diffusion imaging (HARDI). In Wiley Encyclopedia of Electrical and Electronics Engineering (pp. 1–25). John Wiley & Sons, Inc. 10.1002/047134608x.w8258

Dice, L. R. (1945). Measures of the amount of ecologic association between species. Ecology, 26(3), 297–302.

Diedrichsen, J. (2006). A spatially unbiased atlas template of the human cerebellum. NeuroImage, 33(1), 127–138.

Faber, J., Kügler, D., Bahrami, E., Heinz, L.-S., Timmann, D., Ernst, T. M., Deike-Hofmann, K., Klockgether, T., van de Warrenburg, B., van Gaalen, J., Reetz, K., Romanzetti, S., Oz, G., Joers, J. M., Diedrichsen, J., ESMI MRI Study Group, & Reuter, M. (2022). CerebNet: A fast and reliable deep-learning pipeline for detailed cerebellum sub-segmentation. NeuroImage, 264, 119703.

Garyfallidis, E., Brett, M., Amirbekian, B., Rokem, A., van der Walt, S., Descoteaux, M., Nimmo-Smith, I., & Dipy Contributors. (2014). Dipy, a library for the analysis of diffusion MRI data. Frontiers in Neuroinformatics, 8, 8.

Gaviraghi, M., Savini, G., Castellazzi, G., Palesi, F., Rolandi, N., Sacco, S., Pichiecchio, A., Mariani, V., Tartara, E., Tassi, L., Vitali, P., D’Angelo, E., & Wheeler-Kingshott, C. A. M. G. (2021). Automatic Segmentation of Dentate Nuclei for Microstructure Assessment: Example of Application to Temporal Lobe Epilepsy Patients. Computational Diffusion MRI, 263–278.

Glasser, M. F., Sotiropoulos, S. N., Wilson, J. A., Coalson, T. S., Fischl, B., Andersson, J. L., Xu, J., Jbabdi, S., Webster, M., Polimeni, J. R., Van Essen, D. C., Jenkinson, M., & WU-Minn HCP Consortium. (2013). The minimal preprocessing pipelines for the Human Connectome Project. NeuroImage, 80, 105–124.

Hagmann, P., Jonasson, L., Maeder, P., Thiran, J.-P., Wedeen, V. J., & Meuli, R. (2006). Understanding Diffusion MR Imaging Techniques: From Scalar Diffusion-weighted Imaging to Diffusion Tensor Imaging and Beyond. RadioGraphics, 26(suppl_1), S205–S223.

Hancock, D. Y., Fischer, J., Lowe, J. M., Snapp-Childs, W., Pierce, M., Marru, S., Coulter, J. E., Vaughn, M., Beck, B., Merchant, N., Skidmore, E., & Jacobs, G. (2021). Jetstream2: Accelerating cloud computing via Jetstream. *Practice and Experience in Advanced Research Computing*, Article Article 11.

Han, S., Carass, A., He, Y., & Prince, J. L. (2020). Automatic cerebellum anatomical parcellation using U-Net with locally constrained optimization. NeuroImage, 218, 116819.

Herculano-Houzel, S. (2009). The human brain in numbers: a linearly scaled-up primate brain. Frontiers in Human Neuroscience, 3, 31.

Jensen, J. H., & Helpern, J. A. (2010). MRI quantification of non-Gaussian water diffusion by kurtosis analysis. NMR in Biomedicine, 23(7), 698–710.

Jia, J., Staring, M., & Stoel, B. C. (2024). Seg-metrics: a Python package to compute segmentation metrics. In arXiv [cs.CV]. arXiv. http://arxiv.org/abs/2403.07884

Ji, W., Yu, S., Wu, J., Ma, K., Bian, C., Bi, Q., Li, J., Liu, H., Cheng, L., & Zheng, Y. (2021). Learning calibrated medical image segmentation via multi-rater agreement modeling. 2021 *IEEE/CVF Conference on Computer Vision and Pattern Recognition (CVPR)*, 12336–12346.

Jones, D. K. (2004). The effect of gradient sampling schemes on measures derived from diffusion tensor MRI: a Monte Carlo study. Magnetic Resonance in Medicine, 51(4), 807–815.

Jones, D. K., & Basser, P. J. (2004). “Squashing peanuts and smashing pumpkins”: how noise distorts diffusion-weighted MR data. Magnetic Resonance in Medicine, 52(5), 979–993.

Keim, D. A. (1999). Efficient geometry-based similarity search of 3D spatial databases. SIGMOD Record, 28(2), 419–430.

Kim, J., Patriat, R., Kaplan, J., Solomon, O., & Harel, N. (2020). Deep Cerebellar Nuclei Segmentation via Semi-Supervised Deep Context-Aware Learning from 7T Diffusion MRI. *IEEE Access : Practical Innovations*, Open Solutions, 8, 101550–101568.

Kingma, D. P., & Ba, J. (2015). Adam: A Method for Stochastic Optimization. In Y. Bengio & Y. LeCun (Eds.), *3rd International Conference on Learning Representations, ICLR 2015, San Diego, CA, USA, May 7-9*, *2015, Conference Track Proceedings*. http://arxiv.org/abs/1412.6980

Lundell, H., & Steele, C. J. (2024). Cerebellar imaging with diffusion magnetic resonance imaging: approaches, challenges, and potential. Current Opinion in Behavioral Sciences, 56(101353), 101353.

Magin, R. L. (2016). Models of diffusion signal decay in magnetic resonance imaging: Capturing complexity. Concepts in Magnetic Resonance. Part A, Bridging Education and Research, *45A*(4), e21401.

Maier-Hein, L., Reinke, A., Godau, P., Tizabi, M. D., Buettner, F., Christodoulou, E., Glocker, B., Isensee, F., Kleesiek, J., Kozubek, M., Reyes, M., Riegler, M. A., Wiesenfarth, M., Kavur, A. E., Sudre, C. H., Baumgartner, M., Eisenmann, M., Heckmann-Nötzel, D., Rädsch, T., … Jäger, P. F. (2024). Metrics reloaded: recommendations for image analysis validation. Nature Methods, 21(2), 195–212.

Morell-Ortega, S., Ruiz-Perez, M., Gadea, M., Vivo-Hernando, R., Rubio, G., Aparici, F., de la Iglesia-Vaya, M., Catheline, G., Coupé, P., & Manjón, J. V. (2024). DeepCERES: A Deep learning method for cerebellar lobule segmentation using ultra-high resolution multimodal MRI. In arXiv [eess.IV]. arXiv. http://arxiv.org/abs/2401.12074

Murdoch, B. E. (2010). The cerebellum and language: historical perspective and review. Cortex; a Journal Devoted to the Study of the Nervous System and Behavior, 46(7), 858–868.

Myronenko, A. (2019). 3D MRI Brain Tumor Segmentation Using Autoencoder Regularization. *Brainlesion: Glioma, Multiple Sclerosis*, Stroke and Traumatic Brain Injuries, 311–320.

Noguera, C. B., Bao, S., Petersen, K. J., Lopez, A. M., Reid, J. A., Plassard, A. J., Zald, D. H., Claassen, D. O., Dawant, B. M., & Landman, B. A. (2019). Using deep learning for a diffusion-based segmentation of the dentate nucleus and its benefits over atlas-based methods. The Journal of Medical Investigation: JMI, 6(4), 044007.

Riedel, M. C., Ray, K. L., Dick, A. S., Sutherland, M. T., Hernandez, Z., Fox, P. M., Eickhoff, S. B., Fox, P. T., & Laird, A. R. (2015). Meta-analytic connectivity and behavioral parcellation of the human cerebellum. NeuroImage, 117, 327–342.

Semedo, C., Cardoso, M. J., Vos, S. B., Sudre, C. H., Bocchetta, M., Ribbens, A., Smeets, D., Rohrer, J. D., & Ourselin, S. (2018). Thalamic Nuclei Segmentation Using Tractography, Population-Specific Priors and Local Fibre Orientation. Medical Image Computing and Computer Assisted Intervention – MICCAI 2018, 383–391.

Sotiropoulos, S. N., Jbabdi, S., Xu, J., Andersson, J. L., Moeller, S., Auerbach, E. J., Glasser, M. F., Hernandez, M., Sapiro, G., Jenkinson, M., Feinberg, D. A., Yacoub, E., Lenglet, C., Van Essen, D. C., Ugurbil, K., Behrens, T. E. J., & WU-Minn HCP Consortium. (2013). Advances in diffusion MRI acquisition and processing in the Human Connectome Project. NeuroImage, 80, 125–143.

Strick, P. L., Dum, R. P., & Fiez, J. A. (2009). Cerebellum and nonmotor function. Annual Review of Neuroscience, 32, 413–434.

Sun, P., Wu, Y., Chen, G., Wu, J., Shen, D., & Yap, P.-T. (2019). Tissue Segmentation Using Sparse Non-negative Matrix Factorization of Spherical Mean Diffusion MRI Data. Computational Diffusion MRI, 69–76.

Taghanaki, S. A., Zheng, Y., Kevin Zhou, S., Georgescu, B., Sharma, P., Xu, D., Comaniciu, D., & Hamarneh, G. (2019). Combo loss: Handling input and output imbalance in multi-organ segmentation. Computerized Medical Imaging and Graphics: The Official Journal of the Computerized Medical Imaging Society, 75, 24– 33.

Theaud, G., Edde, M., Dumont, M., Zotti, C., Zucchelli, M., Deslauriers-Gauthier, S., Deriche, R., Jodoin, P.-M., & Descoteaux, M. (2022). DORIS: A diffusion MRI-based 10 tissue class deep learning segmentation algorithm tailored to improve anatomically-constrained tractography. Frontiers in Neuroimaging, 1. 10.3389/fnimg.2022.917806

Tournier, J.-D., Calamante, F., & Connelly, A. (2013). Determination of the appropriate b value and number of gradient directions for high-angular-resolution diffusion-weighted imaging. NMR in Biomedicine, 26(12), 1775–1786.

Van Essen, D. C., Smith, S. M., Barch, D. M., Behrens, T. E. J., Yacoub, E., Ugurbil, K., & WU-Minn HCP Consortium. (2013). The WU-Minn Human Connectome Project: an overview. NeuroImage, 80, 62–79.

Virtanen, P., Gommers, R., Oliphant, T. E., Haberland, M., Reddy, T., Cournapeau, D., Burovski, E., Peterson, P., Weckesser, W., Bright, J., van der Walt, S. J., Brett, M., Wilson, J., Millman, K. J., Mayorov, N., Nelson, A. R. J., Jones, E., Kern, R., Larson, E., … SciPy 1.0 Contributors. (2020). SciPy 1.0: fundamental algorithms for scientific computing in Python. Nature Methods, 17(3), 261–272.

Wang, G., Li, W., Aertsen, M., Deprest, J., Ourselin, S., & Vercauteren, T. (2019). Aleatoric uncertainty estimation with test-time augmentation for medical image segmentation with convolutional neural networks. Neurocomputing, 335, 34–45.

Wang, J., Cheng, H., & Newman, S. D. (2020). Sparse representation of DWI images for fully automated brain tissue segmentation. Journal of Neuroscience Methods, 343, 108828.

Woo, B., & Lee, M. (2021, January 31). Comparison of tissue segmentation performance between 2D U-Net and 3D U-Net on brain MR Images. 2021 International Conference on Electronics, Information, and Communication (ICEIC). 2021 International Conference on Electronics, Information, and Communication (ICEIC), Jeju, Korea (South). 10.1109/iceic51217.2021.9369797

Zhang, F., Breger, A., Cho, K. I. K., Ning, L., Westin, C.-F., O’Donnell, L. J., & Pasternak, O. (2021). Deep learning based segmentation of brain tissue from diffusion MRI. NeuroImage, 233, 117934.

Zhang, F., Cho, K. I. K., Seitz-Holland, J., Ning, L., Legarreta, J. H., Rathi, Y., Westin, C.-F., O’Donnell, L. J., & Pasternak, O. (2024). DDParcel: Deep learning anatomical brain parcellation from diffusion MRI. IEEE Transactions on Medical Imaging, 43(3), 1191–1202.

Zhang, L., Tanno, R., Xu, M., Huang, Y., Bronik, K., Jin, C., Jacob, J., Zheng, Y., Shao, L., Ciccarelli, O., Barkhof, F., & Alexander, D. C. (2023). Learning from multiple annotators for medical image segmentation. Pattern Recognition, 138(109400), None. [dataset] Human Connectome Project, WU-Minn Consortium; 2017; 1200 Subjects Data Release (S1200); https://db.humanconnectome.org/

